# A Three-Layered Agent-Based Model of Adult Hippocampal Neurogenesis (HANG-AB3L) with Stochastic Cell Fate Determination

**DOI:** 10.64898/2026.05.08.723711

**Authors:** Pınar Öz, Abdulsamet Atbaşı

## Abstract

Hippocampal adult neurogenesis (HANG) is a highly regulated process where neural stem cells progress through distinct stages—from Type 1 radial glia-like cells to mature neurons—via a complex series of proliferative and differentiative divisions. While recent *in vivo* imaging has provided valuable insights to cellular processes, the exact relationship between individual cell-fate decisions and long-term population stability remains difficult to quantify empirically. In this study, we utilized an agent-based (AB) model to simulate the stochastic dynamics of the hippocampal neurogenic niche. Our results demonstrate that while individual progenitor lineages exhibit high variability and probabilistic division symmetries (proliferative symmetric, asymmetric, and differentiative symmetric), the system achieves deterministic stability as the initial progenitor density increases. We found that the ***T***_**1**_ progenitor pool follows a negative exponential decay profile, with its longevity primarily dictated by the differentiation rate (***α***_***d***,**0**_). Critically, the terminal output of immature neurons (***C***_***IN***,***t***_) was non-linearly coupled to the proliferative capacity of transit-amplifying cells (***α***_***pp***,**0**_); even marginal increases in symmetric proliferative divisions resulted in an exponential expansion of the neuronal pool. These findings suggest that the homeostatic maintenance of the hippocampal niche is governed by a kinetic tuning of division probabilities, providing a theoretical bridge between single-cell stochasticity and robust tissue-level output.

## 1 Introduction

Adult neurogenesis is the remarkable, lifelong process by which newborn neurons are generated from neural stem cells (NSCs) and integrated into existing neural networks (Ming and Song 2011). Located primarily in the subgranular zone (SGZ) of dentate gyrus (DG) and the subventricular zone (SVZ) (Kempermann et al. 2015), the developmental progression of neurogenesis can generally be divided into three distinct phases : First, the division and proliferation of precursor cells; second, the migration and extension of neuronal processes; and third, the structural integration and expression of mature neuronal markers (Kuhn et al. 1996). The progression through these stages is tightly coordinated by post-transcriptional regulation; for example, global protein synthesis undergoes highly dynamic changes, increasing markedly as quiescent NSCs activate to become neurogenic progenitors, dropping significantly during the early neuroblast stage, and increasing again as neurons mature (Baser et al. 2019).

Adult hippocampal neurogenesis (HANG) originates in SGZ (Figure 1), driven by quiescent radial glia-like NSCs, known as Type 1 cells (Kempermann et al. 2015), (Obernier and Alvarez-Buylla 2019). *In vivo* clonal analysis reveals that these individual radial glial precursors are genuinely self-renewing and multipotent, capable of generating both neurons and astrocytes (Bonaguidi et al. 2011). According to the *disposable stem cell* model, once Type 1 cells are activated, they undergo a rapid succession of asymmetric divisions to generate transiently amplifying neural progenitors (Type 2 cells), before eventually undergoing terminal differentiation into mature astrocytes - a process that contributes to the age-related depletion of the NSC pool (Encinas et al. 2011). However, chronic *in vivo* imaging has revealed a more flexible system : while the majority (approx. 80 %) of initial NSC divisions in the SGZ are asymmetric, roughly 14 % are symmetric self-renewing divisions that help expand the stem cell pool (Pilz et al. 2018),(Namba et al. 2011).The resulting Type cells exhibit rapid proliferation and can also undergo symmetric differentiating divisions or asymmetric divisions to significantly expand the clonal lineage (Pilz et al. 2018). These progenitors differentiate into neuroblasts (Type 3 cells), which express immature neuronal markers, such as doublecortin (DCX) and polysialylated neural cell adhesion molecule (PSA-NCAM) (Kempermann et al. 2015). Before complete integration, between 50% and 80% of these young cells undergo apoptosis and are cleared by microglia, serving as a critical quality-control checkpoint (Encinas and Sierra 2012). Surviving neuroblasts migrate a short distance into the inner granule cell layer, extend dendrites into the molecular layer, and project mossy fiber axons to the CA3 region, ultimately maturing into glutamatergic granule cells essential for spatial learning and pattern separation (Ming and Song 2011).

**Fig. 1.**
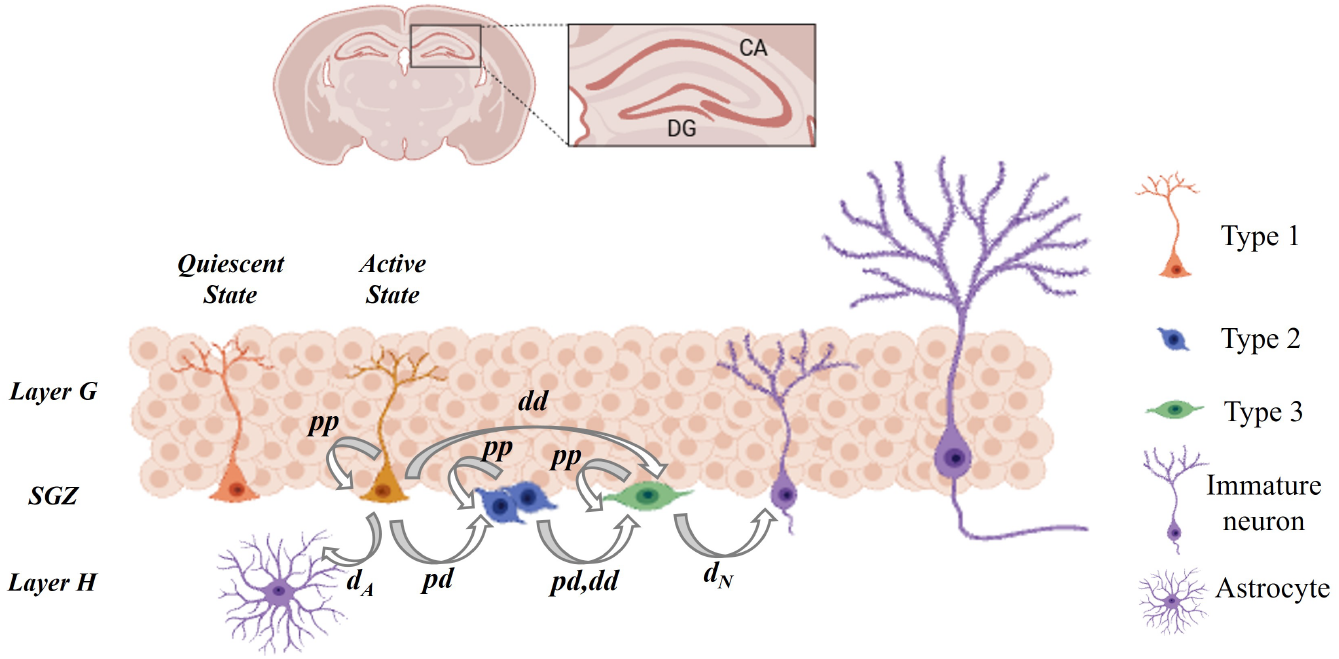
Adult neurogenesis at subgranular zone. The adult neurogenesis in SGZ includes three types of NPCs : Type 1, Type 2 and Type 3. Type 1 NPC can be found either in quiescent state or active state. Once active, Type 1 NPC can either proliferate through symmetric division (*pp*) or differentiate through asymetrical division (*pd*) into Type 2, differentiative symmetric division (*dd*) into Type 3 or division-independent differentiation (*d*_*A*_) into astrocyte. Once differentiated into astrocyte, the cell moves to hilus (Layer H). Type 2 NPCs can either proliferate (pp) or differentiate through asymetrical division (*pd*) or differentiative symmetric division (*dd*) into Type 3. Type 3 NPCs can either proliferate (*pp*) or differentiate through division-independent differentiation (*d*_*N*_) into immature neuron. Once differentiating into immature neuron, the cell migrates to granule cell layer (Layer G). The light-colored cells on the layer G represent adult granule cells. HANG-AB3L model covers the processes until the final differentiation into immature neuron and does not include neuronal maturation.

In contrast to localized integration in the hippocampus, SVZ neurogenesis is characterized by long-distance neuronal migration (Doetsch et al. 1999). SVZ NSCs (Type B cells) share astroglial characteristics with SGZ Type 1 cells (Obernier and Alvarez-Buylla 2019), (Doetsch et al. 1999). Type B cells differentiate into Type C cells through asymmetric division (Obernier et al. 2018). After several rounds of division, Type C cells differentiate into migrating neuroblasts (Type A cells) (Ming and Song 2011), which uniquely assemble into chains and travel extensively along the rostral migratory stream (RMS) toward the olfactory bulb (Ming and Song 2011), (Obernier and Alvarez-Buylla 2019). Once they arrive, they detach from their migratory chains, migrate radially, and terminally differentiate into local GABAergic and dopaminergic interneurons—granule cells and periglomerular cells—that integrate into the olfactory circuitry to support fine odor discrimination (Obernier and Alvarez-Buylla 2019). The mechanisms of NSC proliferation here are highly dependent on symmetric divisions (Obernier et al. 2018). Approximately 20-30% of these cells undergo symmetric self-renewal, while the overwhelming majority (70-80%) undergo symmetric differentiative (or consuming) divisions. This predominant mode generates two transit-amplifying cells per division, effectively decoupling self-renewal from differentiation, which allows massive neuronal production while gradually depleting the stem cell pool over time (Obernier et al. 2018), (Silva-Vargas et al. 2018).

Historically, adult NSCs were assumed to predominantly maintain their populations through invariant asymmetric divisions—yielding one renewed stem cell and one differentiating daughter cell (Silva-Vargas et al. 2018). However, recent clonal lineage-tracing and live-imaging studies have drastically shifted this paradigm, proving that adult NSCs actively undergo symmetric differentiative cell division (also known as symmetric consuming division) (Obernier et al. 2018). In the adult V-SVZ, the vast majority of Type B cells divide symmetrically (Obernier et al. 2018). While about 20% to 30% of these cells undergo symmetric self-renewing divisions (generating two identical stem cells), an overwhelming 70% to 80% undergo symmetric differentiative divisions to generate two transit-amplifying C cells (Pilz et al. 2018), (Obernier et al. 2018). This specific mechanism allows the brain to generate massive numbers of olfactory neurons throughout life while decoupling self-renewal from differentiation, minimizing replicative burden and potential DNA damage in individual stem cells (Obernier et al. 2018). Mathematical modeling based on branching processes in vertebrate neurogenesis further supports this complex balancing act, predicting that the rates of symmetric proliferative, asymmetric, and symmetric differentiative divisions undergo non-monotonic variations to properly orchestrate stem cell maintenance versus neuronal output (Míguez 2015).

Aging profoundly impacts the cellular processes of neurogenesis across both niches. In the hippocampus, chronic *in vivo* imaging reveals that advancing age impairs multiple developmental steps —ranging from the initial cell cycle entry of quiescent NSCs to the survival rate of new cells — ultimately resulting in a drastically reduced clonal output of individual stem cells (Wu et al. 2023). Similarly, in the subependymal zone of aged mice, mathematical modeling and clonal lineage tracing demonstrate altered stem cell behavior. Older stem cell niches exhibit an increased probability of asymmetric stem cell divisions at the expense of symmetric differentiation, coupled with an extended persistence of quiescence between activation phases, driving the age-related decline in overall neurogenesis (Bast et al. 2018).

Computational models of adult neurogenesis translate the extreme biological complexities of stem cell dynamics into measurable mathematical parameters, providing a vital theoretical framework to understand a process that is highly dynamic and challenging to observe entirely *in vivo* (Aimone 2016), (Aimone and Gage 2011). By deliberately focusing on population and cellular dynamics, computational biology utilizes stochastic formalisms, differential equations, and cellular automata to predict the early-stage cellular dynamics of the neurogenic cascade (Aimone 2016). These mathematical models infer biological parameters that are difficult to track experimentally, yielding crucial insights into cell cycle transit times, division symmetry, and stage-specific apoptosis rates (Li et al. 2017). Our study presents an agent-based model to address the cellular dynamics governing the adult hippocampal neurogenesis. Agent-based models enable to track these processes and outcomes at a single cell level and to study population dynamics at the same time. The model focuses solely on the processes starting from the Type 1 NPCs and ending with the final differentiation into immature granule cell.

## 2 Model

Agent-based three-layered adult hippocampal neurogenesis model (HANG-AB3L) is designed on Mesa 2.2.4. package (Ter Hoeven et al. 2025) in Python. The model is based on cellular activities of three neural progenitor cells (NPCs): Radial glia-like Type I cell (*T*_1_), transiently amplifying Type II cell (*T*_2_) and Type III neuroblast (*T*_3_). Each cell type is modeled as an independent neural progenitor agent (NPA), with its own unique and stochastic cellular activity patterns. The model topology includes three independent layers to represent the activities in granule cell layer, SGZ and hilus.

### 2.1 Model Topology

The model topology includes three concentric spherical surfaces to represent the three layers of interest: granule cell layer (Layer G), SGZ and hilus (Layer H).

- **Layer G**: The outermost layer contains mature and immature granule cells. The model assumes all positions are available for an adult-born immature granule cell migrating into this layer from SGZ.
- **SGZ**: The middle layer contains NPC-type agents (NPAs). NPAs are assigned to a
- new location within this layer following the rules of tangential migration, that only occurs at cell birth.
- **Layer H**: The innermost layer contains astrocytes. The model assumes all positions

are available for a newly differentiated astrocyte migrating into this layer from SGZ.

### 2.2 Agent Activity: States, Events and Processes

An agent might start their life as any of the NPAs. The available cellular activity for the agent is determined by the birth event and mother cell type, and follows the neural lineage *T*_1_ →*T*_2_ →*T*_3_. NPAs can be found either in a **quiescence state (***Q*) or an **active state (***A*) (Figure 2). The unique cellular activity of each NPA type includes unique rule sets (Π_*X*_) (§ 2.7) that regulate the probability of six cellular **events**:

**Fig. 2.**
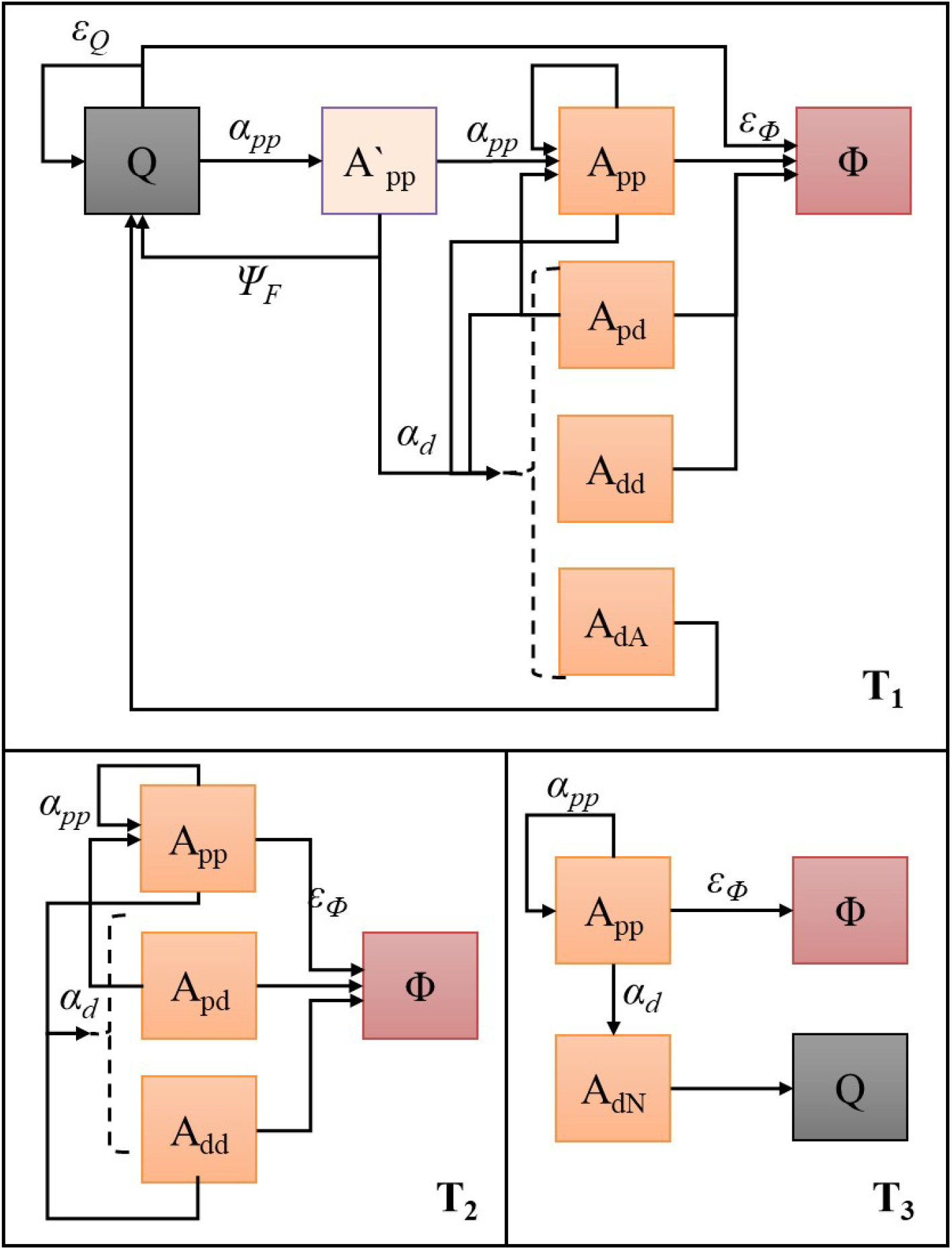
States in HANG-AB3L for each NPA. The states that can follow another state are indicated with arrows. The probability to enter a state is given above the arrow. Light orange box marks the active states (*A*_*pp*_, *A*_*pd*_, *A*_*dd*_) that can follow each other in any order. Dashed line indicates the differentiative states 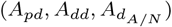 . A *T*_1_ NPA can be found in four different states : quiescent (*Q*), transiently active after the first *pp* event 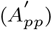, active (*A*_*e*_) where *e* ∈ {*pp, pd, dd, d*_*A*_} and apoptotic (Φ). The NPA can enter 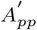 after *Q* and the probability to fall-back to *Q* is determined y Ψ_*F*_ . Φ might also follow *Q*. State *A* can be followed by another *A* or Φ, except 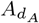 which is followed by *Q. T*_2_ and *T*_3_ NPAs can only be initiated in *A*, which is followed by Φ,except 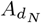, which is followed by *Q*.

1. **Quiescence (***qui*): A dormant, non-dividing state (the phase of the cell cycle) that adult NSCs maintain for extended periods. It is defined as a state (*Q*) with an emission probability *ϵ*_*Q,τ*_ at age *τ* . Only *T*_1_ NPA (*Q*_1_), generated astrocytes (*Q*_*A*_), and immature neurons (*Q*_*N*_) can have quiescence.
2. **Proliferative symmetric cell division (***pp*): The symmetric cell division generates two daughter cells of identical cell type. When a mother cell divides symmetrically to generate a cell of its own type, the event is classified as a proliferative event, as it expands the population of mother cell type. Therefore, in HANG-AB3L, the mother NPA type does not change and a newborn NPA of the same type is generated. The number of proliferative events for an NPA of age *τ* is represented by *ω*_*p,τ*_, and the probability for a proliferative event is represented by *α*_*pp*_.
3. **Differentiative events**: A differentiative event results in at least one agent of a different type from the initial NPA. This includes asymmetric division (*pd*), differentiative symmetric division (*dd*), or division-independent differentiation (*d*_*N*_ or *d*_*A*_).
  - **Asymmetric division (***pd*): It is the major differentiative process in adult neurogenesis (Encinas et al. 2011), (Pilz et al. 2018), (Wu et al. 2023) and results in renewal of the mother cell and generation of a daughter cell of a different type. In HANG-AB3L, the mother NPA type does not change in *pd* and a newborn NPA of another type on the neural lineage is generated.
  - **Differentiative symmetric division (***dd*): symmetric cell division may also result in two identical daughter cells of a type different from the mother.Since there are reports of *dd* in adult neurogenic niche (Pilz et al. 2018),(Obernier et al. 2018), HANG-AB3L includes an event where the mother NPA differentiates along with the production of a different type NPA on the neural lineage.
  - **Differentiation (***d*_*N*_ **or** *d*_*A*_): Differentiation without division is also possible for some NPAs. In HANG-AB3L,the mother NPA itself differentiates into another type without division, which depletes the original NPA type pool. This type of differentiation is allowed only in two instances : The mother NPA differentiates into another agent type on the neural lineage (*d*_*N*_), which is allowed only for *T*_3_ NPAs, and or into an astrocyte (*d*_*A*_), which is only allowed for *T*_1_ NPAs. The number of differentiative events (*pd,dd,d*_*N/A*_) occurred for a NPA until age *τ* is represented by *ω*_*D,τ*_ . The cumulative probability for a differentiative event (*α*_*d*_) is the sum of probability for each differentiative event.
4. **Apoptosis (***ϕ*): Programmed cell death occurs in all NPA types in an age- and activity-dependent manner. The probability of apoptosis is defined as *ϵ*_*ϕ*_ and the state as *ϕ*.

The time after an agent enters the active state is represented with the active age,*τ*_*A*_. State emission probabilities are defined with *ϵ*_*S*_, where 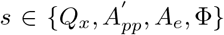 with *x* ∈ {1, *As, IN* } and *e* ∈ {*pp, pd, dd, d*_*N/A*_}. As a general rule,

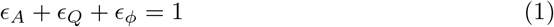

where *ϵ*_*A*_ is the probability to be at an active state.

The probability of each event is determined by cell fate determination mechanics (§ 2.3), that depends on the type, state and age of the NPA, the previously occurred events (internal processes) (§ 2.4) and the niche effects (external processes) (§ 2.5). The occurrence of an event then triggers a series of cellular processes (§ 2.7).

The mother and daughter agent lineages for each state and event is summarized in Table 1.

**Table 1.** Cell lineages with respect to states and events.

| State Class | State at $t = t_{i-1}$ | Probability | Mother Type at $t = t_{i-1}$ | Agent Type at $t = t_i$ | |
| --- | --- | --- | --- | --- | --- |
|  |  |  |  | Mother | Daughter |
| Q | $Q_1$ | $\epsilon_Q$ | $T_1$ | $T_1$ | - |
| | $Q_A$ | 1 | $T_1$ | As | - |
| | $Q_N$ | 1 | $T_3$ | IN | - |
| A | $A_{pp}$ | $\alpha_{pp}$ | $T_1$ | $T_1$ | $T_1$ |
| | | | $T_2$ | $T_2$ | $T_2$ |
| | | | $T_3$ | $T_3$ | $T_3$ |
| | $A_{pd}$ | $\alpha_d$ | $T_1$ | $T_1$ | $T_2$ |
| | | | $T_2$ | $T_2$ | $T_3$ |
| | $A_{dd}$ | $\alpha_d$ | $T_1 / T_2$ | - | $T_3$ |
| | $A_d$ | $\alpha_d$ | $T_1$ | As | - |
| | | | $T_3$ | IN | - |
| $\phi$ | $\phi$ | $\epsilon_\phi$ | $T_1 / T_2 / T_3$ | - | - |

### 2.3 Stochastic Cell Fate Determination Mechanics

The conditional probability function for the processes in the state *s* for agent type *x* is represented by Π_*x*_(*s*) in HANG-AB3L (§ 2.7). Within the condition of Π_*x*_(*s*), cell fate is determined by the random number, *ξ*_*τ*_ ∈ (0, 1). This represents the accumulated stochastic impact of intracellular signal transduction pathways at age *τ*, which are not implemented in the current HANG-AB3L. The variable 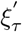 represents the second consecutive *ξ*_*τ*_ called within a single process. Rules of the cell fate determination are then defined over state, event and cell type by comparing *ξ*_*τ*_ with event probability.

### 2.4 Internal Processes

#### 2.4.1 Quiescence Fall-back

*T*_1_ cells can fall back into quiescence (Ψ_*F*_) only after the first few events of proliferation (Encinas et al. 2011). In HANG-AB3L, this fall-back is limited to immediately after a single proliferation event (*ω*_*p*_ = 1). The NPA at *τ*_*A*_ = 1 and *ω*_*P*_ = 1 enters a transitional active state 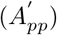, in which the probability for quiescence is reduced by a factor of *κ*_*F*_ .

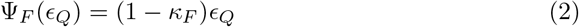

Then, depending on the state of intracellular processes that might impact the fall-back 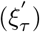, the cell fate is determined as

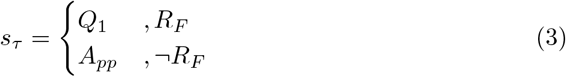

for

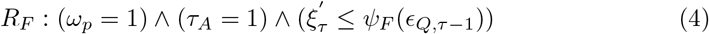

#### 2.4.2 Proliferation Stress

The proliferation stress is defined over the number of proliferative events a NPA goes through and introduced as a stress factor that pushes the cell towards either differentiation or apoptosis, due to the possibility of triggered molecular pathways or telomer shortening. Since these molecular pathways are not implemented in HANG-AB3L, the probability for division is simply reduced by a factor of *κ*_*P*_ by each proliferative event. The proliferative stress can be expressed as a linear system for a NPA at age *τ*, if *s*_*τ*_ = *A*_*pp*_, as

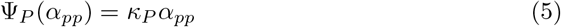

The proliferative stress factor, *κ*_*P*_, is determined with respect to active age, *τ*_*A*_,

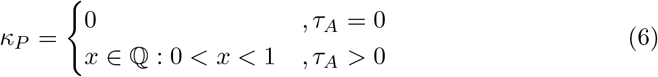

The accumulated effect of proliferative stress can be expressed as

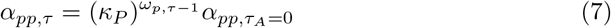

The proliferative stress is also introduced as a limit on the active age (*λ*_1*A*_) for *T*_1_

NPA cellular activity, which acts as a check-point for “differentiate-or-die”.

#### 2.4.3 Differentiation Stress

The internal stress load due to each differentiative event on the mother NPA is introduced as a limit on the number of differentiative events for *T*_1_ and *T*_2_ NPA cellular activity. The number of differentiative events, *ω*_*Dτ*_, is limited with *λ*_*D*1_ for *T*_1_ and *λ*_*D*2_ for *T*_2_ cellular activity.

#### 2.4.4 Aging Stress

The age limit, *λ*_*i*_, for *T*_*i*_ agent, *i* ∈{1, 2, 3}, represent the natural apoptotic processes that might be triggered through several intracellular pathways that are not implemented in HANG-AB3L. The values for *λ*_*i*_ are adjusted with respect to literature (Encinas et al. 2011),(Pilz et al. 2018),(Wu et al. 2023).

### 2.5 External Processes

#### 2.5.1 Crowdedness

Every NPA in SGZ has a “genesis zone” (Figure 3), where the newborn agents move to at their *τ* = 0 and settle down before initiating cellular activity. The spaces in these neighborhoods in SGZ can be previously occupied by a NPA, which means the occupied position is not available for tangential migration. When there is no available space in the genesis zone, a division event goes on with its regular processes, however, can not generate a newborn NPA. This is represented in HANG-AB3L as “stillborn” NPAs, mimicking a reduced form of interactions with neighboring cells that can lead to inhibition of a successful division event. In “stillborn” cases, the event-related processes are carried on, however, the generated NPAs are immediately removed (the event *ex* is immediately triggered) before a placement on the layer. “Stillborn agent” case occurs with the probability *P*_*sb*_ for an NPA **c** with type *x*.

**Fig. 3.**
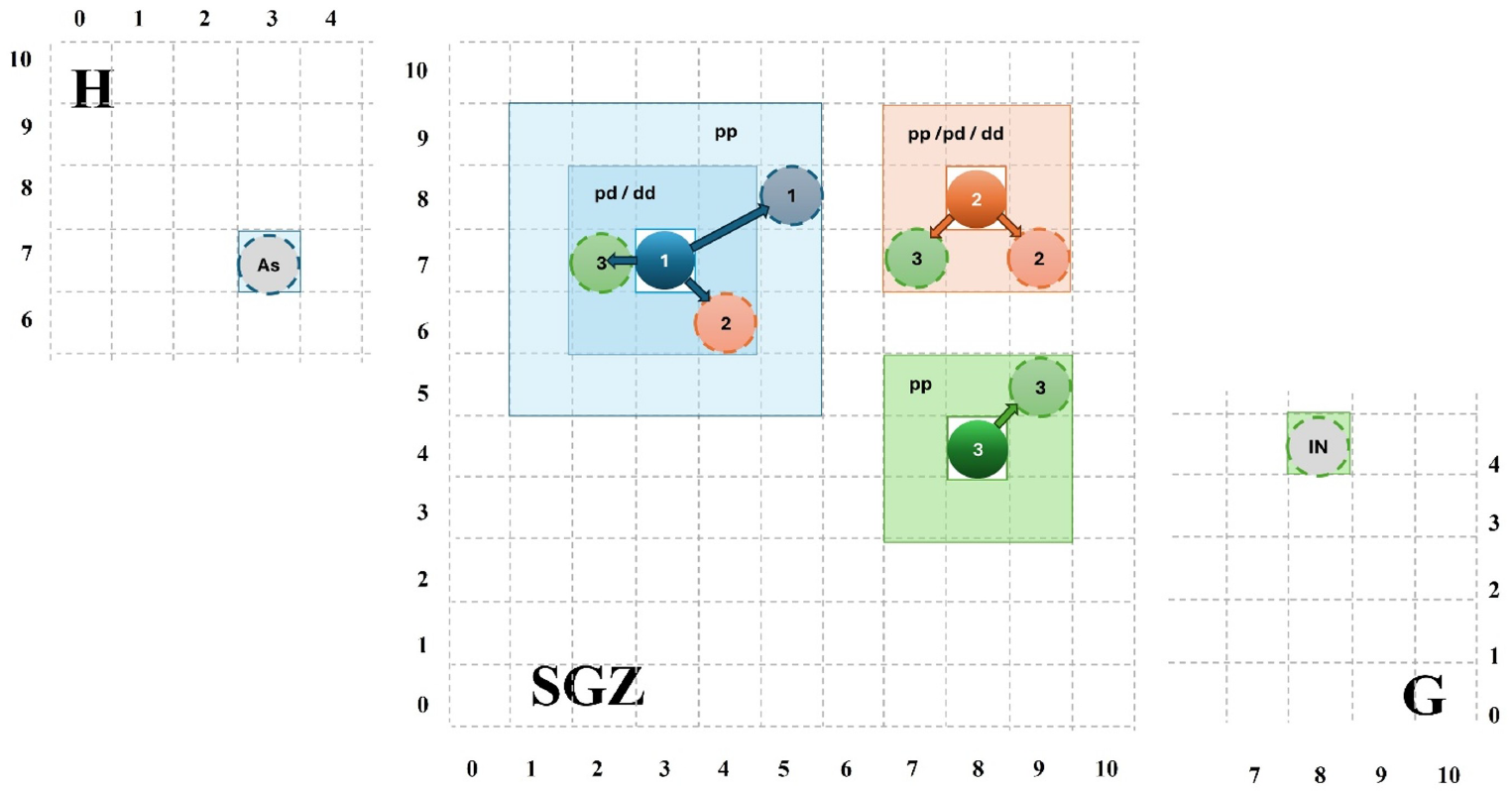
Genesis Zones. In HANG-AB3L, the genesis zones are defined as the areas a newborn NPA can migrate into. A *T*_1_ NPA can produce a newborn *T*_1_ NPA through *pp* that migrates into a position 1-grid away from the mother NPA. Therefore, there are 16 possible positions assuming all positions are available. *T*_2_ and *T*_3_ NPAs can produce newborn NPAs that migrate into adjacent positions, which implies that there are 8 possible positions, assuming all positions are available. The processes *pd* and *dd* produce newborn NPAs that always migrate to adjacent positions. The process *d*_*A*_ results in the migration of differentiated mother NPA to Layer H, while the process *d*_*N*_ results in the migration of differentiated NPA to Layer G. H : Hilus, SGZ : Subgranular zone, G : Granular cell layer. 1 : *T*_1_ agent. Blue areas mark *T*_1_ genesis zone. 2 : *T*_2_ agent. Orange areas mark *T*_2_ genesis zone. 3: *T*_3_ agent. Green areas mark *T*_3_ genesis zone.

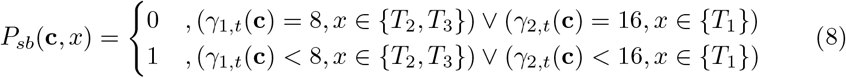

where *γ*_1,*t*_(**c**) give the number of agents adjacent to **c** and *γ*_2,*t*_(**c**) gives the number of agents at 1-grid distance of **c**.

The “stillborn” condition can lead to a compulsory, however, temporary quiescent-like state for all NPAs. Since Ψ_*LI*_ (§ 2.5.2) and Ψ_*P*_ (§ 2.4.2) still applies and the NPA continues to age, long-term crowdedness can push the agent to a “differentiate-or-die” condition. In case of *dd*, both the daughter and the mother agents may die, which may deplete both populations due to long-term crowdedness.

The impact of crowdedness might be exaggerated with lateral inhibition on *T*_1_

NPAs, further depleting the active NPA pool.

#### 2.5.2 Lateral Inhibition

Lateral inhibition on the cellular activity of *T*_1_ cells (Ψ_*LI*_) can be exerted through cell-to-cell Notch signalling (Lampada and Taylor 2023). This molecular pathway is not implemented in HANG-AB3L and reduced to a concept of “crowdedness”.

When a *T*_1_ NPA generates a newborn *T*_1_ NPA through the proliferative event *pp*, the newborn NPA does not “crowd” the adjacent neighborhood and tangentially moves to an empty space 1-grid away from the mother at *τ* = 0. Therefore, the proliferation of a *T*_1_ NPA does not directly lead to an inhibiting crowdedness on itself. However, the proliferation of *T*_1_ NPAs in 2-grid neighborhood can generate newborn *T*_1_ NPAs immediately adjacent to an existing *T*_1_ NPA. Crowdedness in the adjacent neighborhood regulates the proliferative event probability for each *T*_1_ NPA, by attenuating *α*_*pp,τ*_ by a factor of *κ*_*LI*_, which represents the impact of lateral inhibition exerted by its neighbors. The number of adjacent neighbors of a *T*_1_ NPA **c** at a time point *t* is given as *γ*_1,*t*_(**c**). The effect then can be formulated as follows :

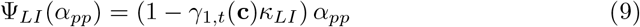

### 2.6 Migration

Any newborn agent moves to a randomly assigned location among the empty positions of their respective “genesis zone” (Figure 3). All NPAs remain at SGZ layer and move by tangential migration. Any agent that differentiates into an astrocyte or an immature neuron move radially to layer H or layer G respectively.

For a *T*_1_ NPA at position [*i, j*], the newborn *T*_1_ NPA tangentially move to an empty position at 1-grid distance after *pp*. The newborn *T*_2_ NPAs after *pd* or newborn *T*_3_ NPAs after *dd* tangentially move to an adjacent empty position. For *dd*, the mother NPA is assumed to remain at [*i, j*] There are 16 available positions for newborn *T*_1_ NPAs and 8 available positions for newborn *T*_2*/*3_ NPAs centered around a *T*_1_ NPA. A *T*_1_ NPA that differentiated into an astrocyte (*d*_*A*_) moves radially to the position [*i, j*] at layer H. Its previous space in SGZ layer becomes empty and available to other NPAs.

For a *T*_2_ NPA at position [*i, j*],the newborn *T*_2_ NPA after *pp* or newborn *T*_3_ NPA after *pd* or *dd* tangentially move to an adjacent empty position. Therefore, there are 8 available positions for newborn *T*_2*/*3_ NPAs centered around a *T*_2_ NPA. For *dd*, the mother NPA is assumed to remain at [*i, j*].

For a *T*_3_ NPA at position [*i, j*], the newborn *T*_3_ NPA after *pp* tangentially move to an adjacent empty position. Therefore, there are 8 available positions for newborn *T*_3_ NPAs centered around a *T*_3_ NPA. A *T*_3_ NPA that differentiated into an immature neuron (*d*_*N*_) moves radially to the position [*i, j*] at layer G. Its previous space in SGZ layer becomes empty and available to other agents.

A dead agent (*ex*) is removed from the map and the space it previously occupied becomes available.

### 2.7 Cellular Processes

The main framework of HANG-AB3L model NPA processes is given in Figure 4.

**Fig. 4.**
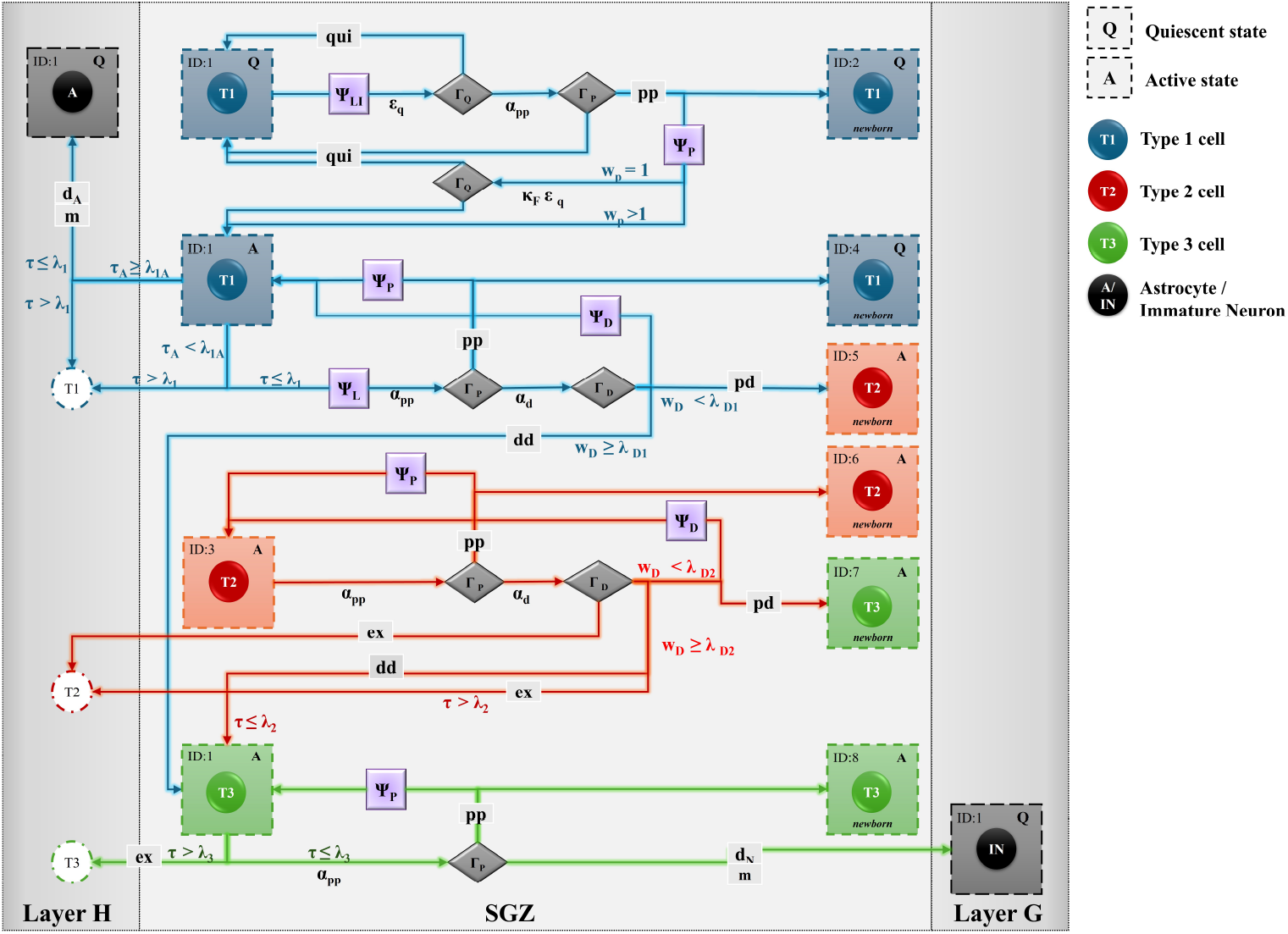
Main framework of HANG-AB3L model NPA processes. The cellular processes and rules for *T*_1_, *T*_2_ and *T*_3_ NPAs are described as the main framework of HANG-AB3L. Processes, states and rule sets of cellular activity are detailed in the respective sections. States are given as square boxes and the NPAs are represented with circles. Circles with dashed lines represent dead NPAs. The possible processes for each NPA is represented with arrows and labeled with the event. The probability or the main governing rule for each event is described above the arrow. Gray diamonds (Γ_*e*_) represent the decision gates for each event, where the rule sets are applied. *Q* : Quiescence state; *A*: Active state; *qui*: quiescence; *pp*: proliferative symmetric division; *pd* : asymmetric division; *dd*: differentiative symmetric division; *d*_*A/N*_ : division-independent differentiation; *ex*: apoptosis; *m*: radial migration; *ϵ*_*Q*_ : probability of quiescence; *α*_*pp*_ : probability of proliferative symmetric division; *α*_*d*_ : probability of differentiative events; *τ* : age of the agent; *τ*_*A*_ : active age of the agent ; *ω*_*P*_ : the number of proliferative events; *ω*_*D*_ : the number of differentiative events; *κ*_*F*_ : coefficient of reduction in quiescence fall-back; *λ*_1*A*_ : maximum active age; *λ*_1*/*2*/*3_ : maximum age for *T*_1*/*2*/*3_ NPAs; *λ*_*D*1*/*2_ : maximum number of differentiative events for *T*_1*/*2_ NPAs; ID: the unique ID of the agent.

#### 2.7.1 *T*_1_ Cellular Processes

*T*_1_ NPA cellular activity is age-dependent and activity-dependent. A *T*_1_ NPA can remain in *Q*_1_ as long as it is not activated with another event (*τ >* 0,*τ*_*A*_ = 0). Newborn *T*_1_ NPA starts at *Q*_1_ (*ϵ*_*Q,τ*=0_ = 1) with *ω*_*D,τ*=0_ = 0,*ω*_*P,τ*=0_ = 0. After first day, the probability for remaining in *Q*_1_ becomes

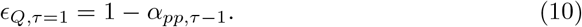

If a *T*_1_ NPA falls back to quiescence at age *τ*, then *ϵ*_*Q,τ*_ = *ϵ*_*Q,τ*=1_ (§ 2.4.1).

The proliferative event probability, *α*_*pp,τ*_, is regulated with both Ψ_*LI*_ and Ψ_*P*_ for *T*_1_ NPA. As a rule, lateral inhibition, Ψ_*LI*_, always preceeds the proliferative stress, Ψ_*P*_ . Since there are only two modulatory factors on *α*_*pp*_, the change in *α*_*pp,τ*_ for a *T*_1_ agent at age *τ* is dependent on the number of neighboring *T*_1_ NPAs and the state of the NPA at *τ* − 1, which can be expressed as

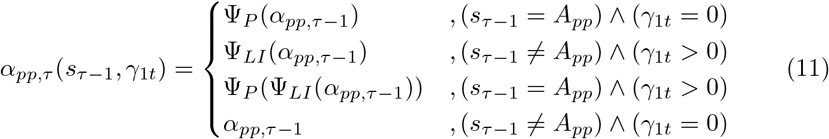

The inhibition is presumed to be lasting long enough that removal of an inhibiting NPA does not immediately release the inhibition, representing the lingering intracellular processes. Therefore, together with Ψ_*P*_, Ψ_*LI*_ can push a *T*_1_ NPA to a compulsory quiescence.

Recent findings suggest that *T*_1_ cells can symmetrically divide into two *T*_3_ cells (Pilz et al. 2018),(Obernier et al. 2018). Therefore, our model incorporates *dd* into *T*_3_.

The *T*_1_ NPA cellular activity is expressed with Π_1_(*s*_*τ*_) as follows :

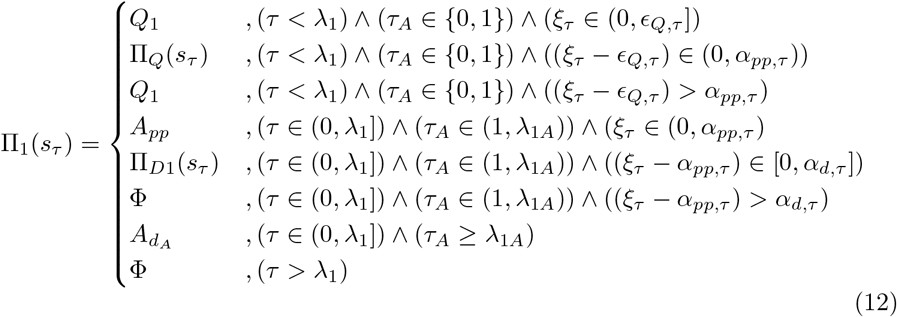

Where

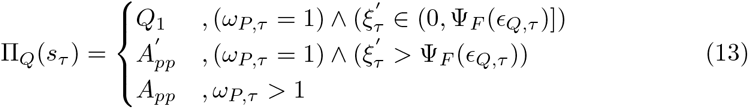

And

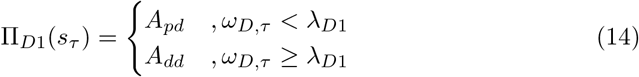

Therefore, the change in the *T*_1_ NPA population size at *t*, **C**_1,*t*_, is

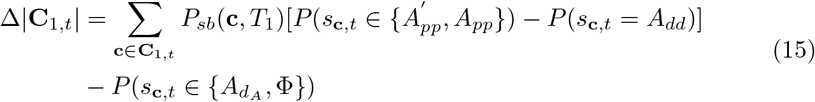

where *P* is the indicator function. For the agent **c**, *x*_*c*_ is the type of **c** and *s*_**c**,*t*_ is the state of **c** at time *t*.

The change in number of permanently quiescent agents in **C**_1,*t*_ at time *t* (*N*_*QP,t*_) can be estimated as

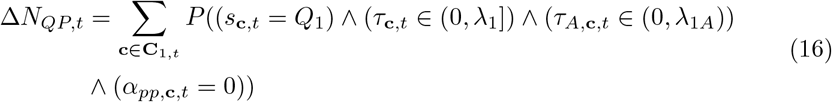

with *N*_*QP,t*=0_ = 0.

The change in the size of **C**_*As,t*_ is dependent only on the activity of NPAs in **C**_1,*t*_.

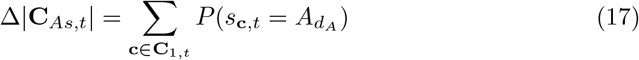

#### 2.7.2 *T*_2_ Cellular Processes

*T*_2_ cellular activity is age-dependent. A newborn *T*_2_ NPA at *τ* = 0 starts at active state (*τ*_*A*_ = 1) and *ϵ*_Φ,*τ*=0_ = 0. However, there is no activity immediately after birth. The proliferative event limit, *α*_*pp,τ*_, for *T*_2_ NPA is regulated only with Ψ_*P*_, therefore,

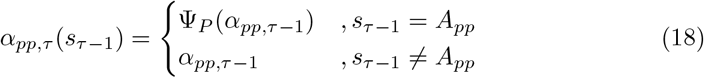

The *T*_2_ NPA cellular activity is expressed with Π_2_(*s*_*τ*_) as follows :

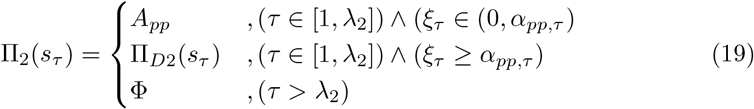

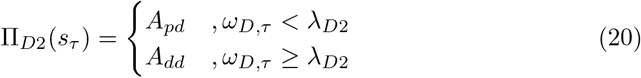

The change in the *T*_2_ NPA population size at *t*,**C**_2,*t*_, is

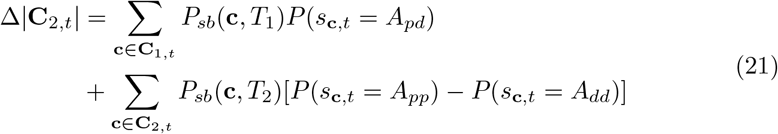

#### 2.7.3 *T*_3_ Cellular Processes

*T*_3_ NPA cellular activity is age-dependent. A newborn *T*_3_ NPA at *τ* = 0 starts at active state (*τ*_*A*_ = 1). In the cases of *dd* or *d*_*N*_, where the mother NPA has differentiated and aged, there is a possibility that the *T*_3_ NPA may die before any activity if 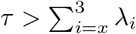 for mother NPA *T*_*x*_, *x* ∈ {1, 2}.

The proliferative event limit, *α*_*pp,τ*_, for *T*_3_ NPA is not regulated with Ψ_*P*_ or any internal/external process. The *T*_3_ NPA cellular activity is expressed with Π_3_(*s*_*τ*_) as follows :

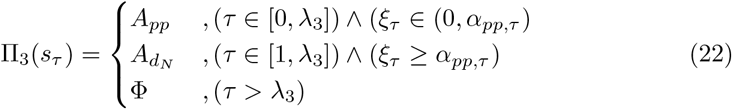

Instead of *λ*_3_, the age limit for a *T*_3_ NPA is *λ*_2_ + *λ*_3_ if it differentiated from a *T*_2_ NPA and *λ*_1_ + *λ*_3_ if it differentiated from a *T*_1_ NPA.

The change in the size of **C**_3,*t*_ is dependent on the activity of NPAs of **C**_1,*t*_,**C**_2,*t*_ and **C**_3,*t*_.

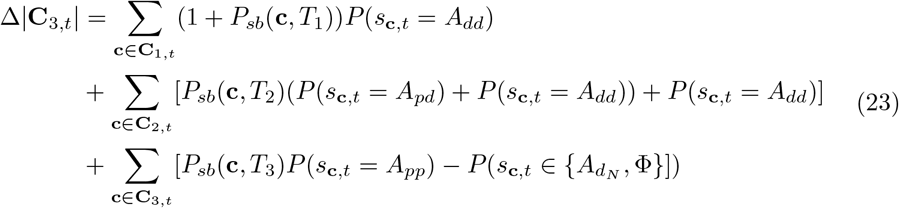

The change in the size of **C**_*IN,t*_ is dependent only on the activity of NPAs in **C**_3,*t*_.

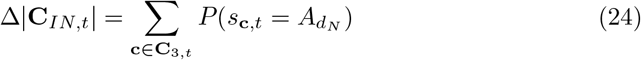

### 2.8 Parameter Selection, Visualization and Simulation

Initial parameters were selected with respect to existing literature (Bonaguidi et al. 2011; Encinas et al. 2011; Pilz et al. 2018; Wu et al. 2023) as given in Table 2. For the initial *T*_1_ NPAs at day 0, *α*_*pp*,0_ + *α*_*d*,0_ + *ϵ*_*Q*,0_ = 1. *α*_*pp*,0_ and *α*_*d*,0_ were initiated with 0.1 and altered by a step of 0.1 in parametric tests. The simulation was run for a minimum of 100 days with a time step of 1 day. In parametric tests, the map was set to 100 x 100 grids. The initial *T*_1_ NPA location was randomized among trials. In lineage tracking, two *T*_1_ NPA were initiated on a 10×10 grid, where the same initial parameters were used.

**Table 2.** Default initial parameters in HANG-AB3L model.

| Parameter | Initial Value |
| --- | --- |
| Differentiation limit for $T_1$ NPA ($\lambda_{D1}$) | 10 |
| Differentiation limit for $T_2$ NPA ($\lambda_{D2}$) | 5 |
| Active age limit for $T_1$ NPA ($\lambda_{1A}$) | 10 |
| Maximum age limit for $T_1$ NPA ($\lambda_1$) | 30 |
| Maximum age limit for $T_2$ NPA ($\lambda_2$) | 5 |
| Maximum age limit for $T_3$ NPA ($\lambda_3$) | 5 |
| Coefficient of quiescence fall-back ($\kappa_F$) | 0.75 |
| Coefficient of proliferation stress ($\kappa_P$) | 0.6 |
| Coefficient of crowdedness ($\kappa_{LI}$) | 0.125 |

For the GUI visualization of HANG-AB3L, inherent visualization tools of Mesa (v2.2.4) package was utilized (Supplementary Video 1). The parametric tests were run for 100 days with 100 repetitions and the population growth and parametric curves were obtained as average and standard deviation from repeated tests.

## 3 Numerical Results

### 3.1 NPA Lineages in HANG-AGB3L

A single NPA lineage can be virtually tracked in HANG-AB3L. As an example, two *T*_1_ NPAs were randomly located on a 10×10 grid environment for lineage tracking (Figure 5-A). The initial parameters were set as in Table 2, with *α*_*pp*,0_ = *α*_*d*,0_ = 0.3. In this example, one of the initial NPAs remained quiescent for 21 days and died 1 day after the first *pp* event. The newborn *T*_1_ NPA survived and remained quiescent for 13 days. The second initial *T*_1_ NPA went through 1 *pp* and 1 *pd* event before *ex* (apoptosis). However, this was enough for the generation of new *T*_2_ and *T*_3_ NPAs, ending with the generation of 2 newborn immature neurons (Figure 5-B). The first immature neuron was differentiated on 10^*th*^ day and the second on 12^*th*^ day. At 41^*st*^ day, all NPAs were depleted. This lineage did not include the rare *dd* event; all differentiation events were through either *pd* (for *T*_1_ and *T*_2_ NPAs) or *d*_*N*_ (for *T*_3_ NPAs). It should be noted that, due to the stochastic nature of cell fate determination, every new simulation generates NPAs with different lineages.

**Fig. 5.**
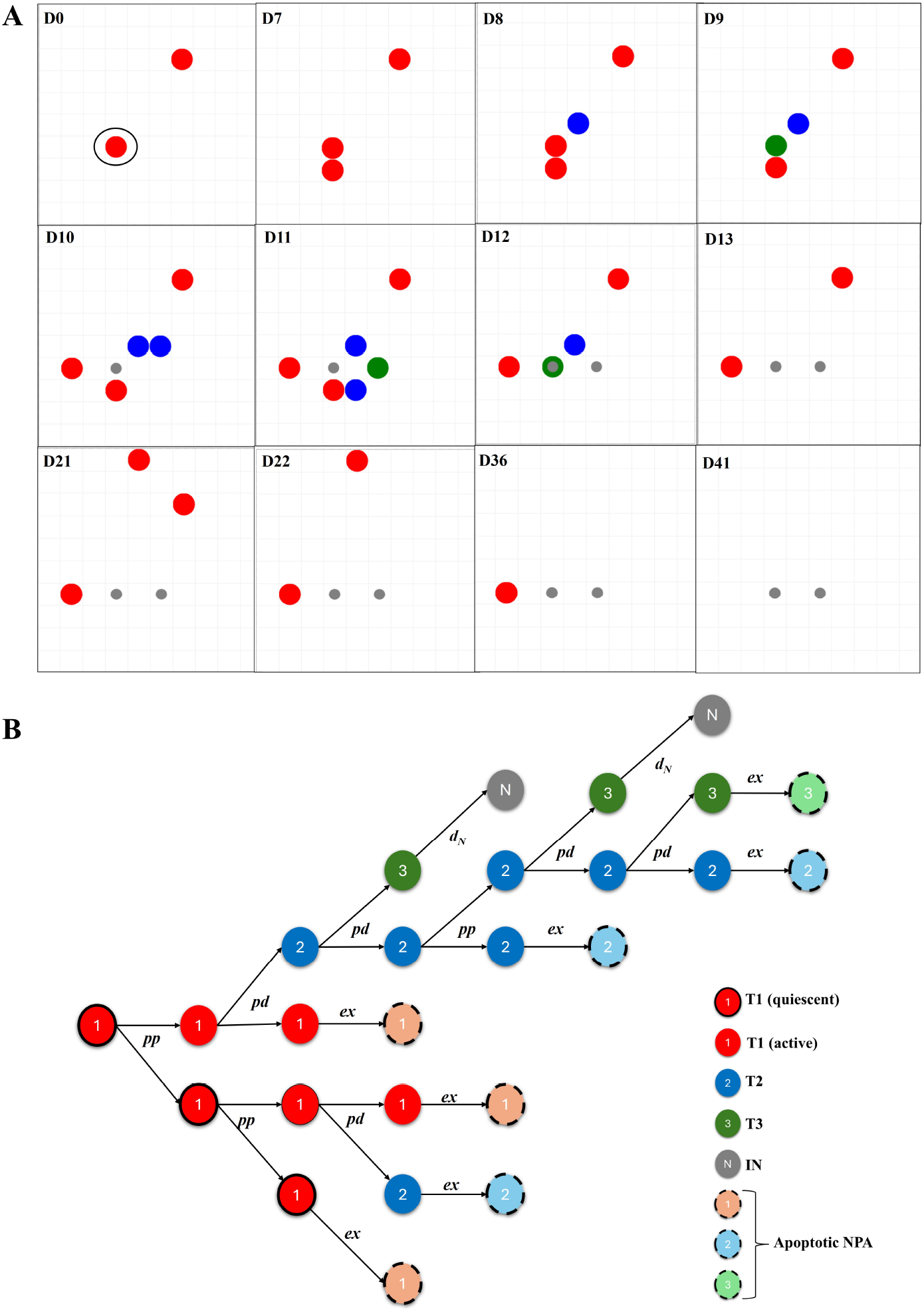
NPA Lineages. **(A)** The population was initiated with 2 *T*_1_ NPAs randomly located on a 10×10 grid at day 0 (D0). The events were tracked until all NPAs are depleted. Red: *T*_1_ NPA, Blue: *T*_2_ NPA, Green: *T*_3_ NPA, Gray : Immature Neuron. All NPAs are in the same layer (SGZ), whereas the immature neurons are on an upper (visually overlapping) layer. **(B)** The lineage of circled *T*_1_ NPA at D0 in **(A)** is tracked through generations and events. *pp* : Proliferative symmetric division, *pd*: Asymmetric division, *d*_*N*_ : Division-independent differentiation in neural lineage, *ex* : Apoptosis, IN : Immature neuron.

### 3.2 *T*_1_ NPA Quiescence and NPA Population Growth in Crowded Environments

The influence of initial NPA density (*N*) on population stability was assessed across a 5, 100, 400 and 800 initial *T*_1_ NPAs (Figure 6). At sparse environment (*N* = 5) (Figure 7), the system exhibited significant stochastic fluctuations, highlighting the impact of individual events on total population variance. As *N* increased, the stochastic trajectories converged towards the deterministic model and the impact of random drift due to stochastic cell fate diminished. At *N* ≥ 400 (Figure 11 and 13), the system followed predictable, first order kinetics, consistent with the law of large numbers in biological modelling.

**Fig. 6.**
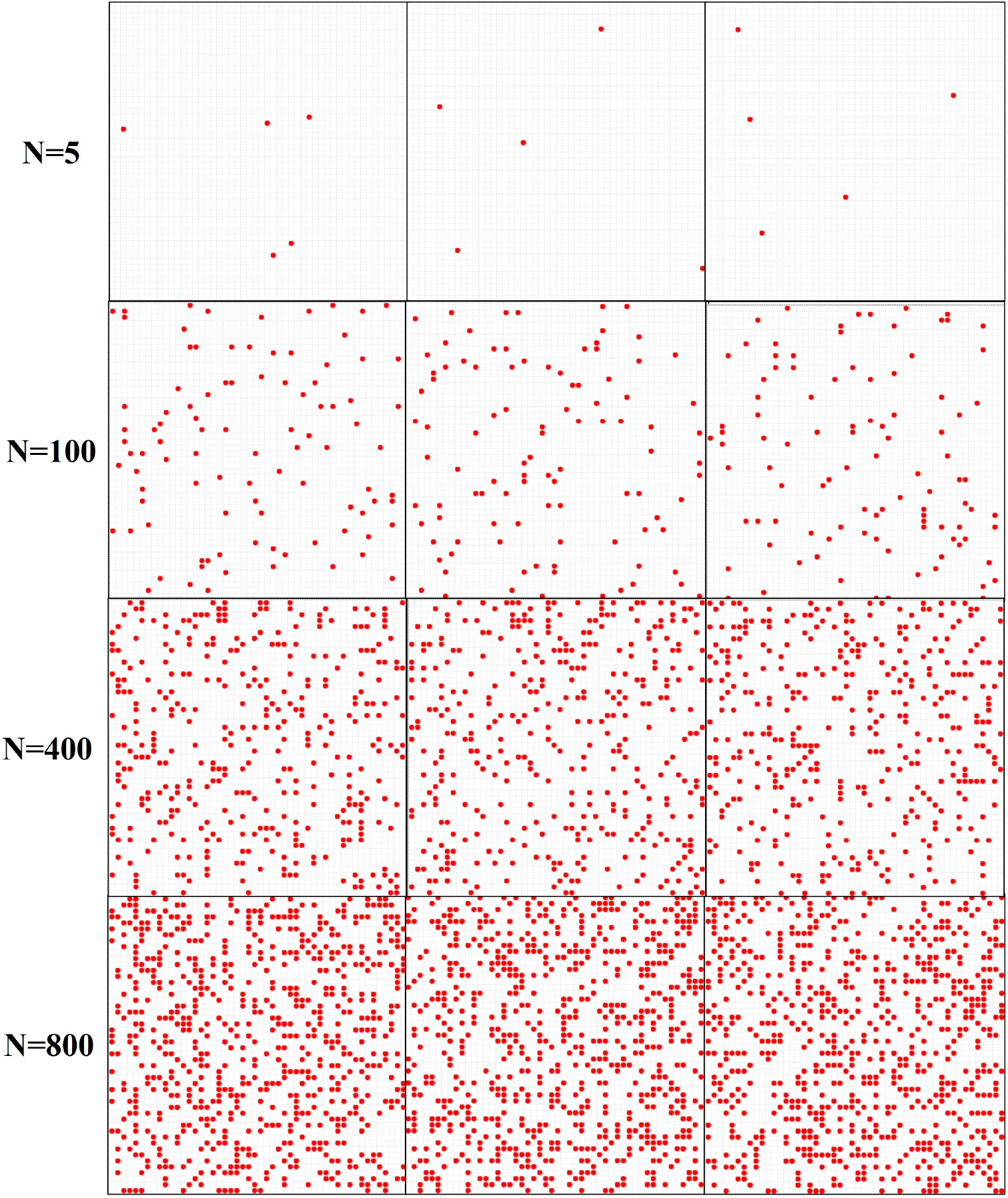
Sparse and Crowded NPA Environments. HANG-AB3L model was simualted for the initial *T*_1_ NPA population sizes 5, 100, 400 and 800 on a 100×100 grid environment. Red circles represent *T*_1_ NPAs. All *T*_1_ NPAs were initiated at quiescent state and were randomly located within the environment for each run. Three examples for the initial positions are given in the figure.

**Fig. 7.**
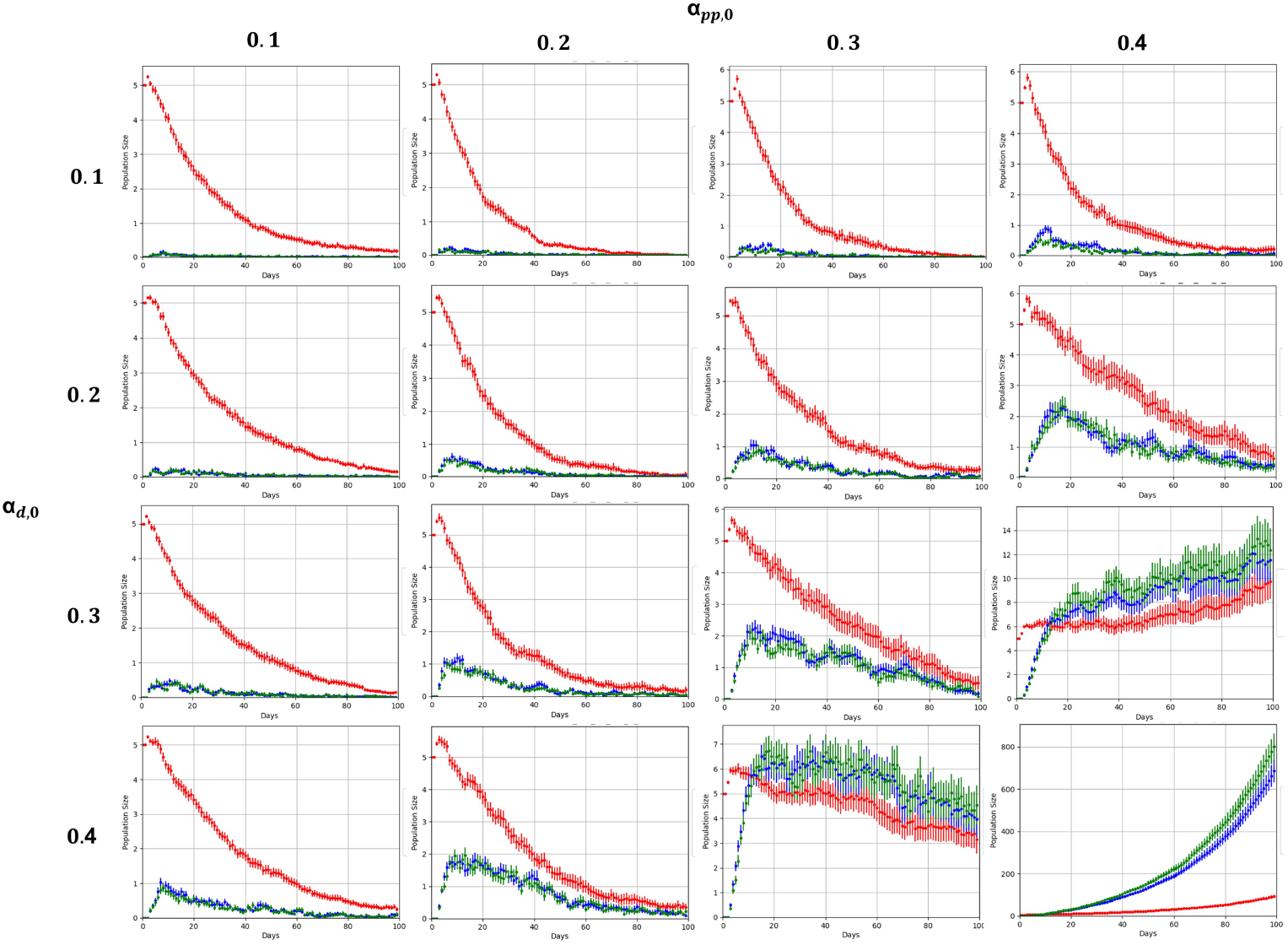
NPA Population Growth with 5 *T*_1_ NPA at *t* = 0. The population growth for *T*_1_ (red),*T*_2_ (blue) and *T*_3_ (green) NPA populations are given for varying *α*_*pp*,0_ and *α*_*d*,0_ as temporal averages and standard deviation over 100 repetitions.

Across all *N*, the *T*_1_ NPA population consistently followed a negative exponential decay profile following a transient peak (approximately at 4-8 days) (Figure 16) and extent of the decay is determined by *α*_*pp*,0_ and *α*_*d*,0_. The time to attenuate to half of the initial population size (*t*_1*/*2_) of the *T*_1_ NPA population was inversely proportional to the *α*_*d*,0_. Increasing *α*_*d*,0_ from 0.1 to 0.4 resulted in a marked reduction of *T*_1_ quiescence, accelerating the depletion of the progenitor pool and shifting the system toward a lower-steady equilibrium. At lower *α*_*d*,0_ values (0.1) and higher *N* (Figure 11,13 and 15), the *T*_1_ population maintained a significant reservoir even after 100 days.

The initial *N* was positively correlated with the level of quiescent plateau in *T*_1_ NPA population (Figure 15). For *N* = 100 (Figure 9), quiescent reservoir was only maintained for *α*_*d*,0_ = 0.1 at approximately 40% of *N* at *t* ≥ 40 days. For *N* = 400 (Figure 11), a quiescent reservoir was preserved at approximately 75% at *t* ≥ 40 days for *α*_*d*,0_ = 0.1, 30% at *t* ≥ 55 days for *α*_*d*,0_ = 0.2 and 12.5% at *t* ≥ 60 days for *α*_*d*,0_ = 0.3.Across all *α*_*d*,0_, a quiescent reservoir was preserved for *N* = 800 (Figure 13) at approximately 87.5% at *t* ≥ 40 for *α*_*d*,0_ = 0.1, 31% at *t* ≥ 60 days for *α*_*d*,0_ = 0.3 and 12.5% at *t* ≥ 60 days for *α*_*d*,0_ = 0.4 . For *α*_*d*,0_ = 0.2, *α*_*pp*,0_ ≥ 0.3 increased the quiescent proportion from approximately 50% to 62.5% and pulled the time to reach the plateau from 70 to 60 days.

The transition of *α*_*pp*,0_ from 0.1 to 0.4 demonstrates a nonlinear increase in downstream NPA yield in all *N* (Figure 7,9,11 and 13). While *T*_1_ NPA decay remains relatively constant (Figure 15), the *T*_2_ (Figure 16)/ *T*_3_ NPA (Figure 17) peaks scale dramatically . This confirms *α*_*pp*,0_ as the primary driver of clonal expansion within these populations.

The downstream populations (*T*_2_ and *T*_3_ NPAs) exhibited transiently amplifying characteristics of coupled source-sink systems (Figure 16 and 17). For all *N, T*_2_ and *T*_3_ NPA populations displayed nearly identical temporal profiles, suggesting a rapid, high fidelity transition between these states with minimal residence time in the intermediate *T*_2_ NPA phase. The magnitude of *T*_2_/*T*_3_ NPA burst was synergistically controlled by both *α*_*pp*,0_ and *α*_*d*,0_. While *α*_*d*,0_ governed the rate of entry into the differentiated pool, higher values of *α*_*pp*,0_ (0.3-0.4) significantly increased the peak amplitude and shifted the maximum population density toward later time points, which was still within the first 10 days for both *T*_2_ NPA (Figure 16) and *T*_3_ NPA (Figure 17).

Under conditions of low initial *α*_*pp*,0_ and *α*_*d*,0_, the system maintained a prolonged *T*_1_ NPA presence with minimal downstream flux. Conversely, high-rate regimes favored a rapid, high-magnitude pulse of differentiation, resulting in the eventual exhaustion of the progenitor pool in favor of a transient surge in differentiated *T*_2_/*T*_3_ NPAs. *T*_1_ NPA acts as the primary regulator of system longevity, its decay rate determines the duration of the differentiation pulse. *T*_2_/*T*_3_ NPAs act as the functional output of the system, their magnitude is decoupled from *T*_1_ NPA longevity and is instead governed by the proliferative coefficient. The system transitions from a stochastic regime at low NPA numbers to a highly predictable deterministic regime at higher densities, ensuring robust lineage output despite the probabilistic nature of individual cellular decisions.

### 3.3 Neuronal Differentiation in Sparse and Crowded Environments

The immature neuron populations across all *N* was positively correlated with *α*_*pp*,0_, *α*_*d*,0_ and N (Figure 8,10,12 and 14) and followed a trend that can be described as 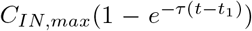, where *t*_1_ was approximately 4-5 days in all cases. The accumulation of immature neurons follows a saturating growth model where the final population size is limited by the initial NPA pools but scaled exponentially by the proliferative capacity of *T*_2_/*T*_3_ NPAs. The system reaches a stable homeostatic plateau by 60–80th day across all *N* . *α*_*pp*,0_ is the most sensitive parameter for controlling total neuronal output, outweighing the impact of *N* .

**Fig. 8.**
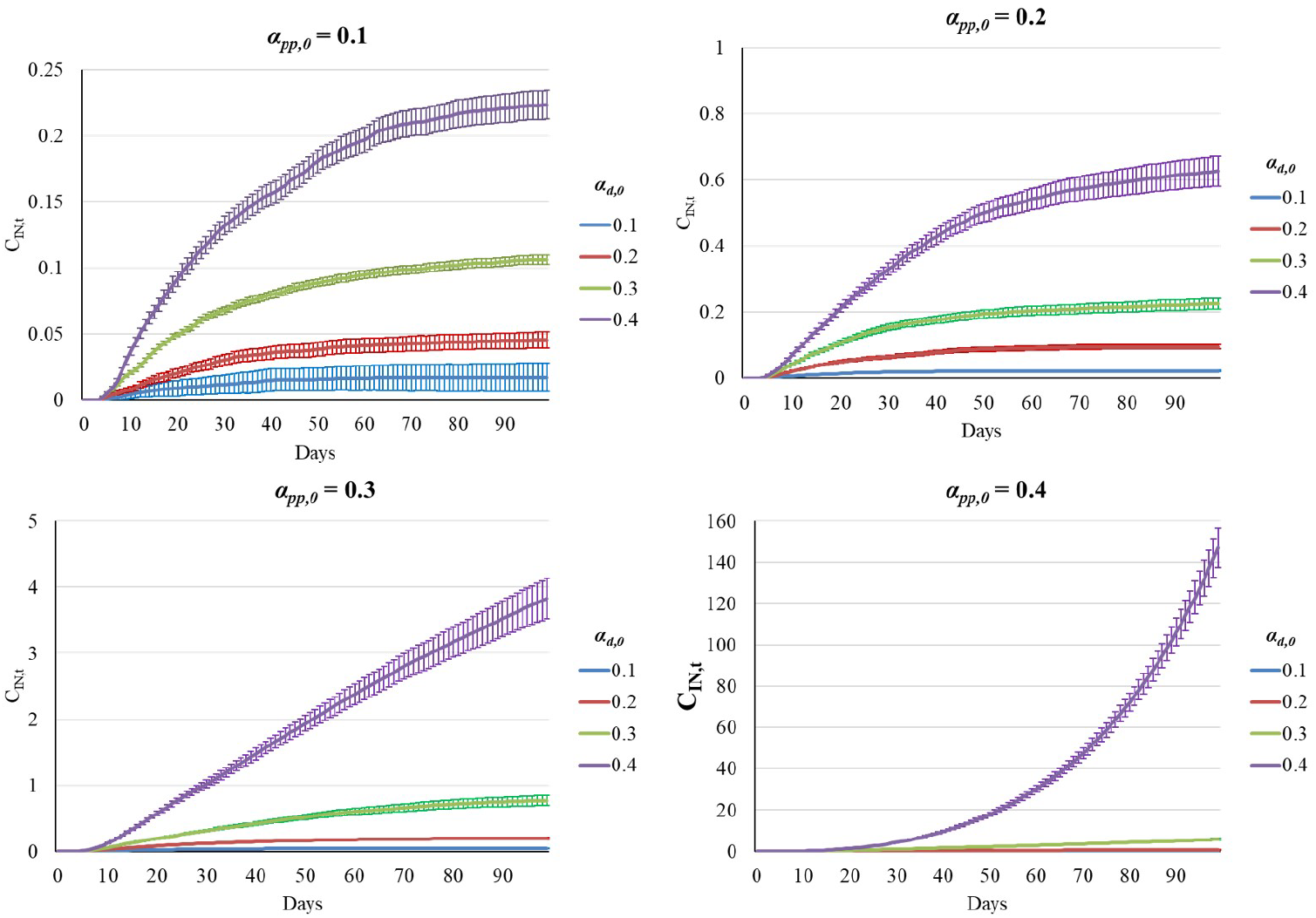
Immature Neuron Population Growth with 5 *T*_1_ NPA at *t* = 0. The immature neuron population growth (C_*IN,t*_) are given for varying *α*_*pp*,0_ and *α*_*d*,0_ as temporal averages and standard deviation over 100 repetitions.

**Fig. 9.**
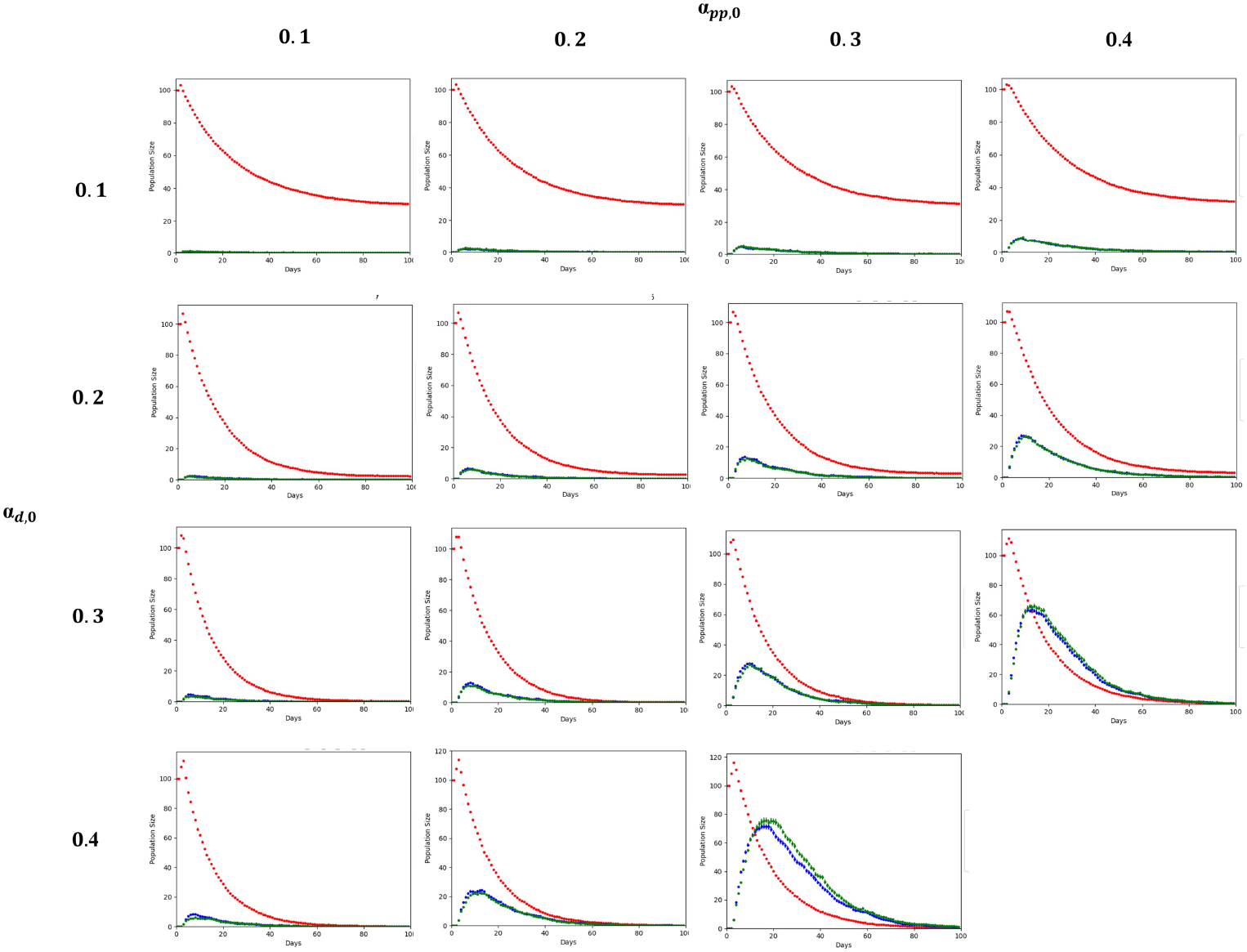
NPA Population Growth with 100 *T*_1_ NPA at *t* = 0. The population growth for *T*_1_ (red),*T*_2_ (blue) and *T*_3_ (green) NPA populations are given for varying *α*_*pp*,0_ and *α*_*d*,0_ as temporal averages and standard deviation over 100 repetitions.

As demonstrated in Supplementary Figure 1, all division types exhibit a transient impulse response, peaking within the first 10–20 days before undergoing exponential decay, a profile that mirrors the depletion of the primary *T*_1_ NPA pool. The ratio of these division events is critically sensitive to the *α*_*pp*,0_ and *α*_*d*,0_.Analysis of the division fraction (*n*_*pp*_*/n*_*total*_) in Supplementary Figure 2 reveals that higher *α*_*d*,0_ suppress the relative frequency of *pp*, effectively forcing the system toward terminal differentiation. Conversely, increasing *α*_*pp*,0_ stabilizes the *n*_*pp*_ fraction at a higher steady-state baseline (approximately 0.8 in the *α*_*pp*,0_ = 0.4 regime), maintaining a robust NPA reservoir. This relationship is further quantified by the proliferation/differentiation ratio (Supplementary Figure 3), where low differentiation rates coupled with high proliferative capacity yield a high-ratio environment characterized by significant stochastic noise. The cumulative impact of these division dynamics is manifested in the immature neuron (**C**_*IN,t*_) accumulation curves (Figure 8,10,12 and 14). The transition from *N* = 5 to *N* = 400 demonstrates that while early-stage stochasticity in division choice can cause order-of-magnitude variance in small populations, large-scale systems achieve a stable, sigmoidal accumulation pattern. The final neuronal yield is non-linearly coupled to the proliferation/differentiation ratio: even a marginal increase in the *pp* frequency early in the lineage results in an exponential expansion of the terminal **C**_*IN,t*_ pool. Collectively, these data suggest that the neurogenic output of the system is not merely a function of initial NPA numbers, but is primarily governed by the kinetic tuning of cell division events for especially *T*_2_/*T*_3_ NPAs.

**Fig. 10.**
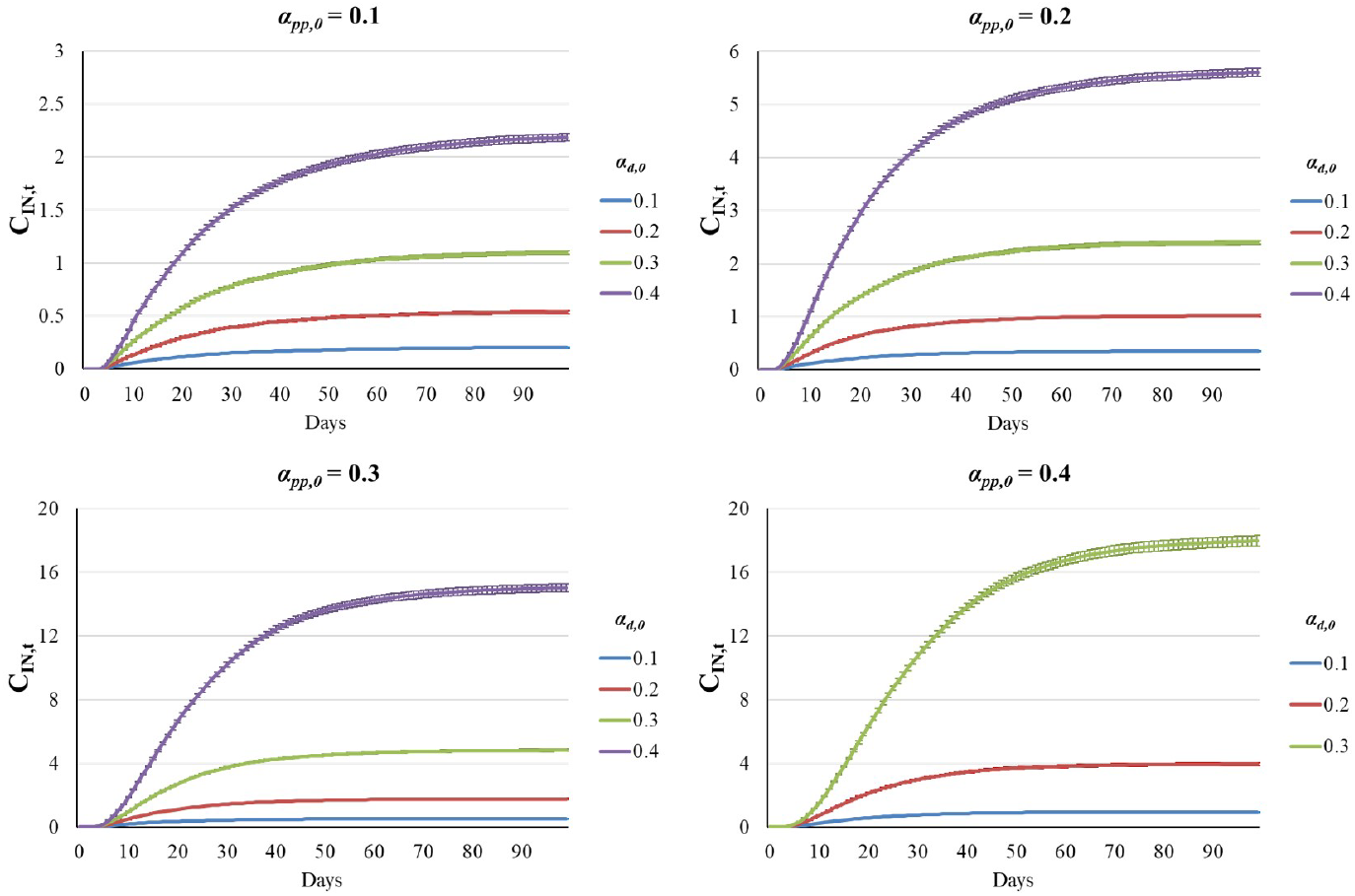
Immature Neuron Population Growth with 100 *T*_1_ NPA at *t* = 0. The immature neuron population growth (C_*IN,t*_) are given for varying *α*_*pp*,0_ and *α*_*d*,0_ as temporal averages and standard deviation over 100 repetitions.

**Fig. 11.**
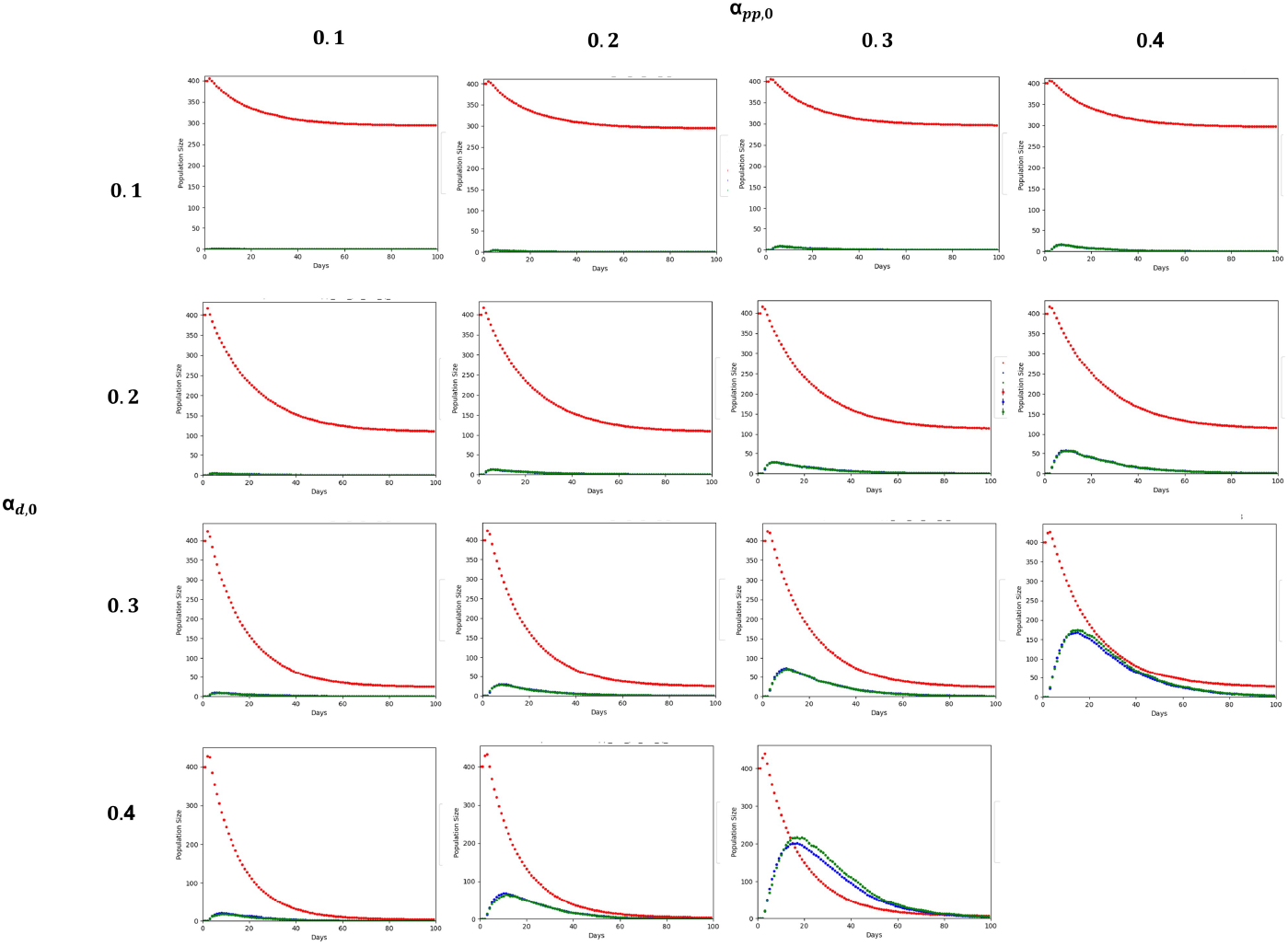
NPA Population Growth with 400 *T*_1_ NPA at *t* = 0. The population growth for *T*_1_ (red),*T*_2_ (blue) and *T*_3_ (green) NPA populations are given for varying *α*_*pp*,0_ and *α*_*d*,0_ as temporal averages and standard deviation over 100 repetitions.

**Fig. 12.**
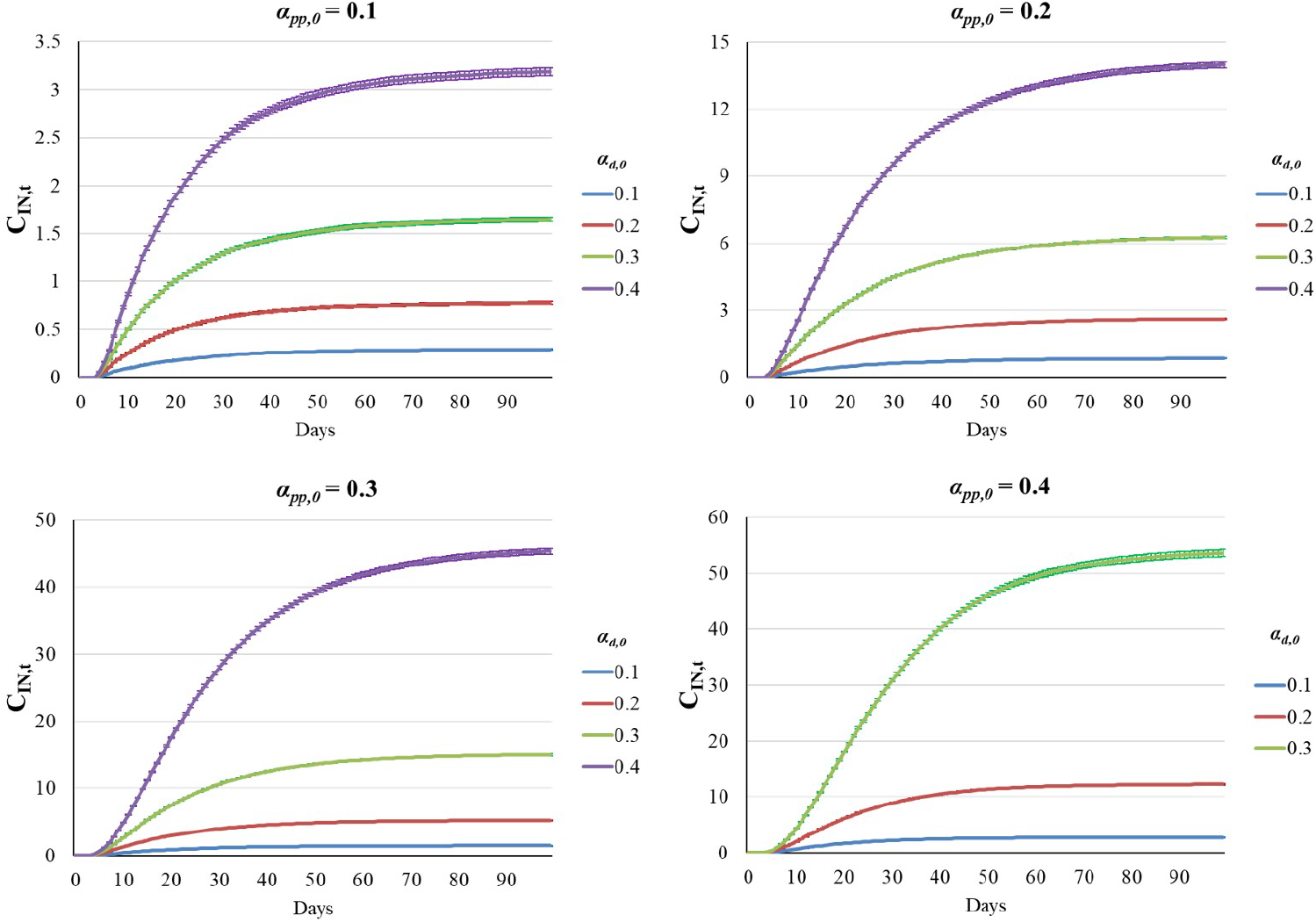
Immature Neuron Population Growth with 400 *T*_1_ NPA at *t* = 0. The immature neuron population growth (C_*IN,t*_) are given for varying *α*_*pp*,0_ and *α*_*d*,0_ as temporal averages and standard deviation over 100 repetitions.

**Fig. 13.**
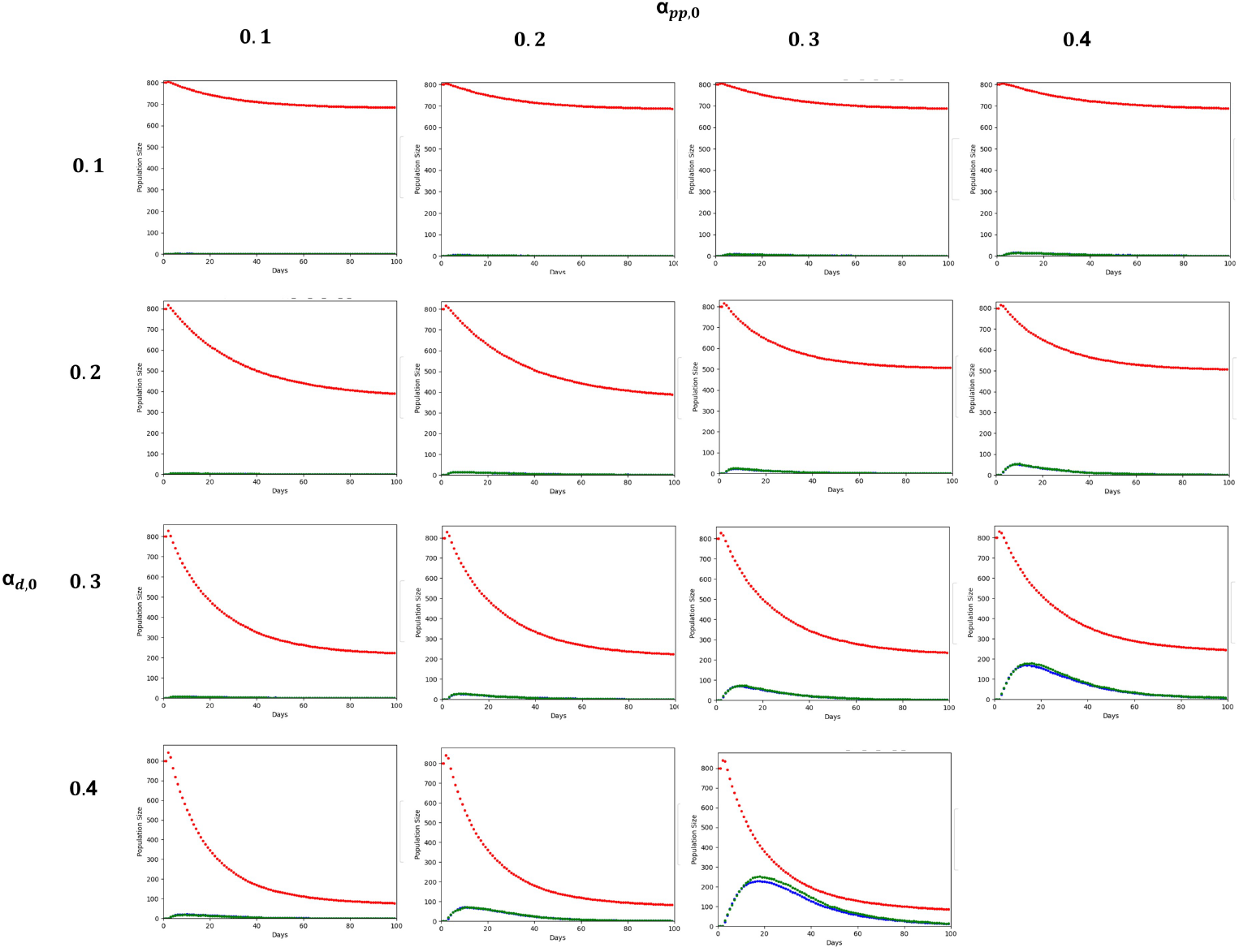
NPA Population Growth with 800 *T*_1_ NPA at *t* = 0. The population growth for *T*_1_ (red),*T*_2_ (blue) and *T*_3_ (green) NPA populations are given for varying *α*_*pp*,0_ and *α*_*d*,0_ as temporal averages and standard deviation over 100 repetitions.

**Fig. 14.**
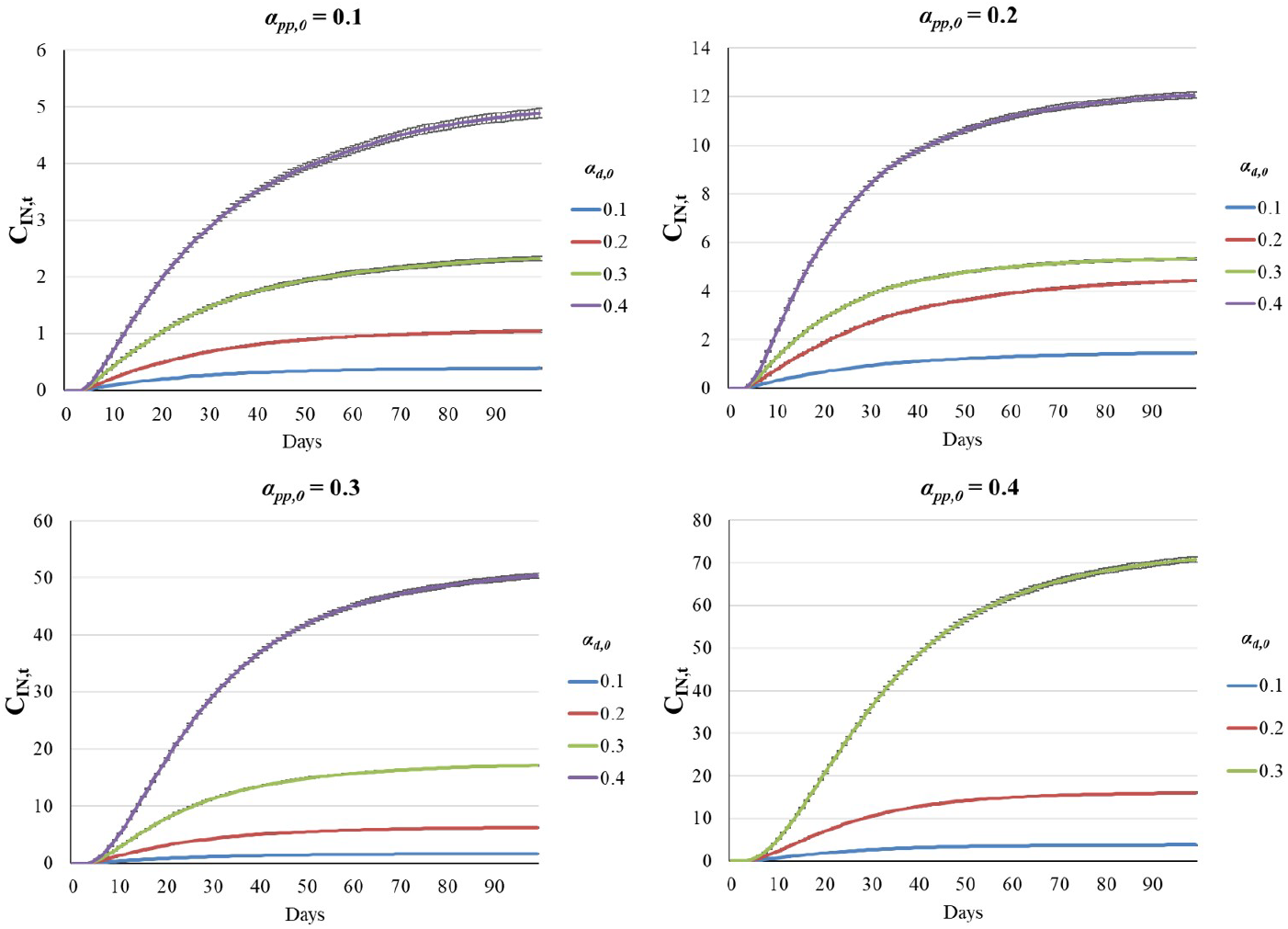
Immature Neuron Population Growth with 800 *T*_1_ NPA at *t* = 0. The immature neuron population growth (C_*IN,t*_) are given for varying *α*_*pp*,0_ and *α*_*d*,0_ as temporal averages and standard deviation over 100 repetitions.

**Fig. 15.**
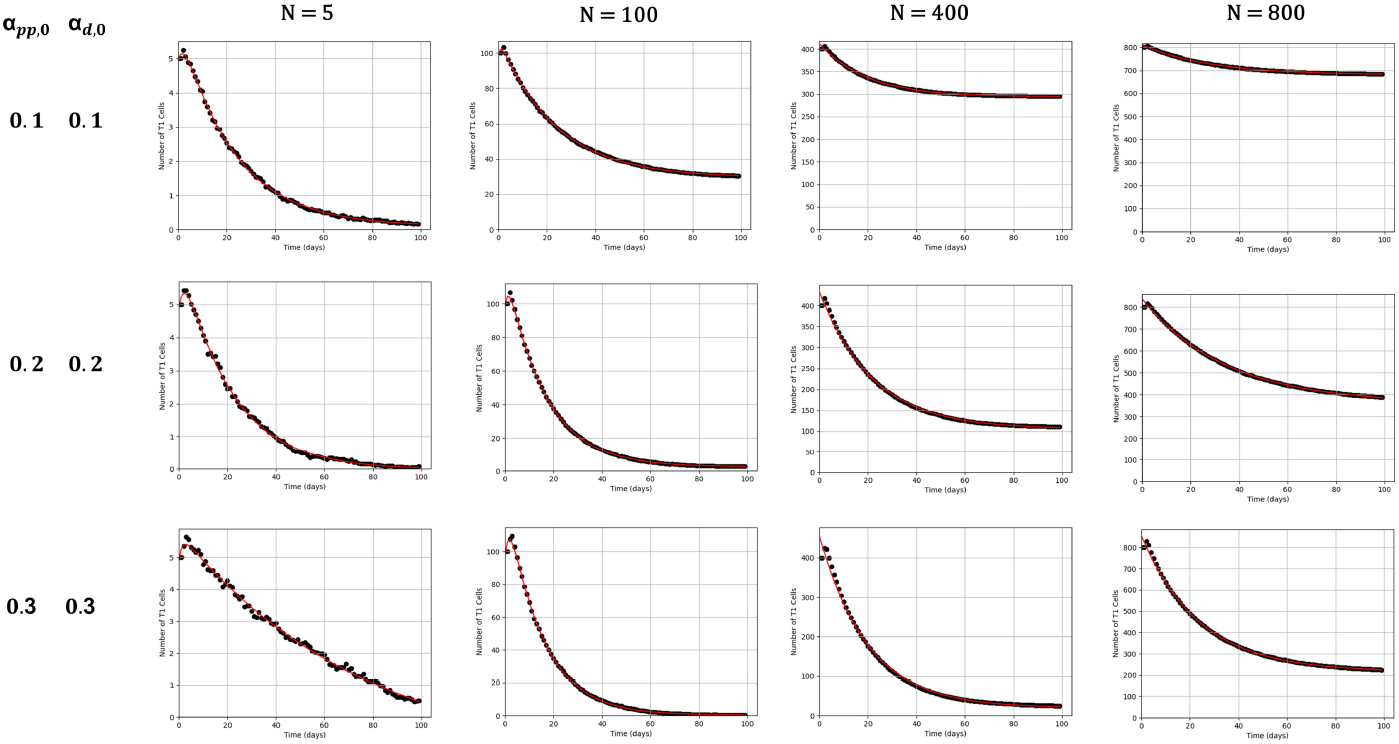
*T*_1_ NPA Population Growth for Sparse and Crowded Environments. *T*_1_ NPA population growth (black dot) was given for varying *N, α*_*pp*,0_ and *α*_*d*,0_ with fitting curve (red line) described with the equation 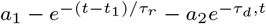 .

**Fig. 16.**
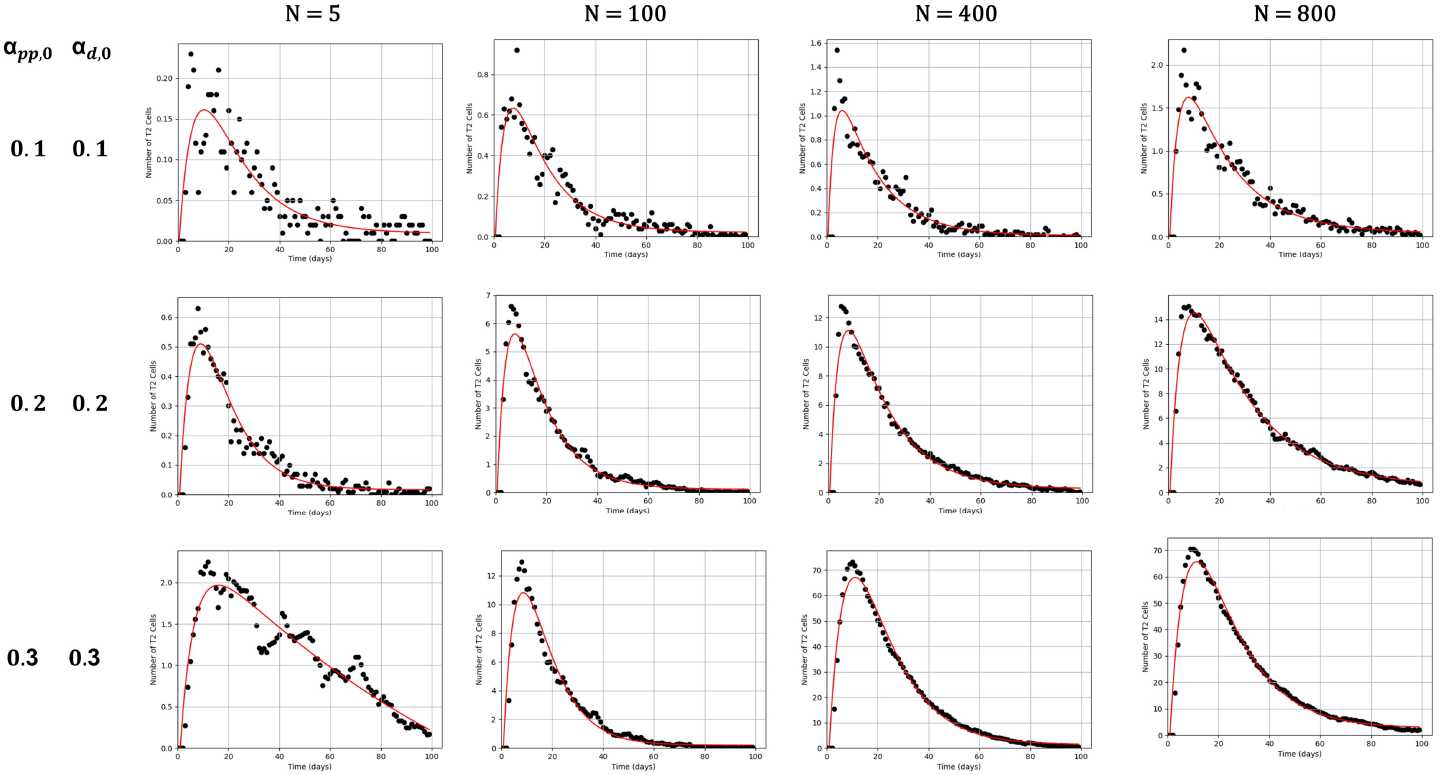
*T*_2_ NPA Population Growth for Sparse and Crowded Environments. *T*_1_ NPA population growth (black dot) was given for varying *N, α*_*pp*,0_ and *α*_*d*,0_ with fitting curve (red line) described with the equation 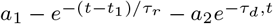 .

**Fig. 17.**
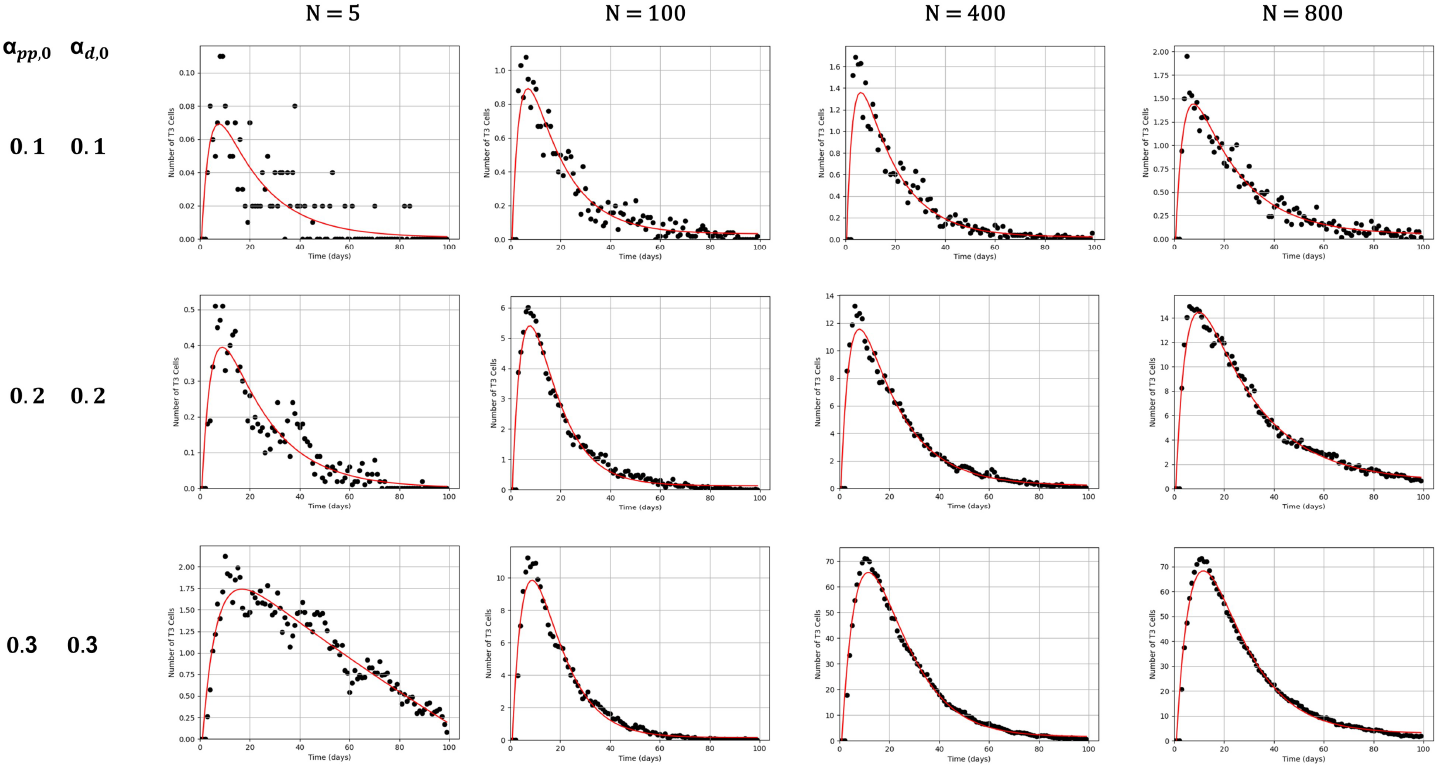
*T*_3_ NPA Population Growth for Sparse and Crowded Environments. *T*_1_ NPA population growth (black dot) was given for varying *N, α*_*pp*,0_ and *α*_*d*,0_ with fitting curve (red line) described with the equation 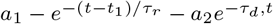 .

## 4 Discussion

Computational models of adult neurogenesis may employ a wide range of methods. To model the early stages of neurogenesis and the stochastic nature of cell fate decisions, researchers employed branching process formalisms (Barton et al. 2014). Cell lineage tree models utilize binary trees to track the derivation of neurons from progenitor cells over successive generations (Slater et al. 2009). By applying metrics such as Colless’s Index to measure the symmetry of these cell lineage trees, models have demonstrated that stochastic, generation-dependent probabilities can accurately reproduce the patterns of mammalian neurogenesis (Slater et al. 2009). The Multitype Bellman-Harris branching model has been used to estimate exact apoptosis rates and cell-cycle transit times, predicting that cell death is highest specifically during the neuroblast stage, and that the transit-amplifying progenitor (TAP) renewal probability is remarkably low (Li et al. 2017). Other branching models detail how the dynamics of stem cell differentiation are characterized by non-monotonic variations in the rates of symmetric versus asymmetric divisions to properly balance population maintenance (Míguez 2015). Deterministic models using ordinary differential equations (ODEs) and partial differential equations (PDEs) are highly effective for simulating large-scale population dynamics and physical cell migration. Multiscale mathematical models employing PDEs have successfully tracked the progression of intermediate progenitors through specific cell cycle phases (G1, S, G2, M) over developmental time, providing accurate estimations of cell kinetics and neuronal output in the cerebral cortex (Postel et al. 2019). In the SVZ, PDE models predict the mechanisms of cell migration, capturing how cells diffuse, proliferate, differentiate, and undergo apoptosis (Van Schepdael et al. 2013). Furthermore, ODE models have been vital in evaluating age-related changes in HANG (Ziebell et al. 2014). By fitting these models to *invivo* data, researchers have predicted that the age-related decline is driven by altered stem cell dynamics; specifically, older stem cells exhibit an extended persistence of quiescence and an increasing probability of asymmetric stem cell divisions rather than symmetric differentiation (Bast et al. 2018), (Ziebell et al. 2018). To explore spatial dynamics and single-cell interactions, cellular automata (CA) models operate on discrete grids where rules governing proliferation, migration, differentiation, and death are applied to individual agents (Lehotzky and Zupanc 2019). CA models evaluate how developmental processes are influenced by local factors and can simulate tissue-level phenomena, such as the growth of neurospheres and contact inhibition, emerging purely from stochastic behavioral variations at the single-cell level (Lehotzky and Zupanc 2019). Finally, other stochastic single-cell models have investigated the molecular coupling between cell cycle progression and differentiation (Stopka and Boareto 2019). These models reveal that a coupled regulation of cell cycle and differentiation restricts population overgrowth within a stem cell population (Stopka and Boareto 2019).

The results from HANG-AB3L highlight the critical interplay between stochastic cell-fate decisions and deterministic population outcomes in HANG. Our findings demonstrate that while individual progenitor lineages exhibit high variability—characterized by sporadic quiescence and probabilistic division symmetries (*pp, pd, dd*) — the population behavior follows predictable first-order kinetics as the initial progenitor density (*N*) increases. This convergence aligns with broader computational frameworks, where stochastic formalisms and branching process models are utilized to reconcile single-cell behaviors with global lineage patterns. Specifically, our observation that the *T*_1_ NPA pool follows a negative exponential decay profile, with longevity regulated by the differentiation rate (*α*_*d*,0_), mirrors previous ODE models (Ö z 2019) suggesting that age-related neurogenic decline is driven by altered stem cell quiescence and a shift toward asymmetric division.

Furthermore, the burst dynamics observed in the transiently-amplifying (*T*_2_/*T*_3_ NPA) populations underscore the role of *α*_*pp*,0_ as the primary determinant of terminal neuronal yield. The high-fidelity coupling between *T*_2_ and *T*_3_ NPA states observed here reflects multiscale PDE models that track rapid progression through intermediate cell cycle phases. Interestingly, our results indicate that even marginal shifts in the proliferation /differentiation ratio early in the lineage can exponentially scale the immature neuron (**C**_*IN,t*_) pool. This supports the molecular coupling theories which suggest that tightly regulated cell cycle and differentiation rates are necessary to prevent population overgrowth while ensuring robust lineage output. By integrating these agent-based spatial interactions with kinetic tuning, HANG-AB3L provides a theoretical bridge between the probabilistic nature of local cell-fate decisions and the homeostatic maintenance of the HANG niche.

HANG-AB3L model also demonstrates a more realistic system behavior, especially for highly crowded environment, due to the crowdedness and Ψ_*LI*_ processes implemented in the model. The model might be expanded by the additional intracellular and extracellular processes that regulate HANG, such as extracellular BDNF and GABA concentrations. Overall, HANG-AB3L provides a robust and highly adaptive tool to study the cellular processes in HANG in a controlled simulation environment.

## List of Abbreviations

A: Active state
As: Astrocyte
c: An agent
C_*t*_: The population size at time *t*
*dd*: Differentiative symmetric cell division
*d*_*N*_: Division-independent differentiation into immature neuron
*d*_*A*_: Division-independent differentiation into astrocyte
IN: Immature neuron
NPA: Neural progenitor agent
*P*: The indicator function
*P*_*sb*_: The probability of a stillborn agent due to the effect of crowdedness
*pp*: Proliferative symmetric cell division
*pd*: Asymmetric cell division
*Q*: Quiescent state
*s*_*τ*_: The state of an agent at age *τ*
*T*_1_: Radial glia-like cells (Type I)
*T*_2_: Transiently amplifying cells (Type II)
*T*_3_: Neuroblasts (Type III)
*α*_*pp*_: The probability for proliferative events
*α*_*d*_: The probability for differentiative events
*γ*_*t*_: The number of neighboring cells at time *t*
*ϵ*_*A*_: The emission probability for being at an active state
*ϵ*_*Q*_: The emission probability for being at a quiescence state
*ϵ*_Φ_: The emission probability for being at an apoptotic state
*κ*_*F*_: Coefficient of quiescence fall-back
*κ*_*P*_: Coefficient of proliferation stress
*κ*_*LI*_: Coefficient of crowdedness
*λ*_1*A*_: The maximum active age
*λ*_*D*_: The maximum number of differentiative events
*λ*_*i*_: The maximum age for NPA of type *T*_*i*_
*ξ*_*τ*_: Random number representing the accumulated stochastic impact of intracellular signal transduction pathways at age *τ*
Π_*x*_: Cellular activity rule sets for a cell of type *x*
*τ*: Age of a NPA
*τ*_*A*_: Active age of a NPA
Φ: Apoptosis state
Ψ_*F*_: The process of quiescence fall-back
Ψ_*P*_: The process of proliferation stress
Ψ_*LI*_: The process of lateral inhibition
*ω*_*p,τ*_: The number of *pp* events at age *τ*
*ω*_*D,τ*_: The number of differentiative events at age *τ*

## Supplementary information

**Supplementary Figure 1 : Event counts with 800** *T*_1_ **NPA at t=0**. Event counts (black dot) was given for varying *α*_*pp*,0_ and *α*_*d*,0_ with fitting curve (red line) described with the equation 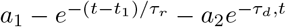.

**Supplementary Figure 2: Ratio of** *pp* **events to the total count of cell divisions with 800** *T*_1_ **NPAs at t=0**. The number of proliferative symmetric divisions (*pp*) to total division events was regulated by the *α*_*pp*_ and *α*_*d*_.

**Supplementary Figure 3 : Ratio of proliferation to differentiation with 800** *T*_1_ **NPAs at t=0**. Ratio of proliferative divisions (*pp*) to differentiative divisions (*pd,dd*) on each day were regulated by the *α*_*pp*_ and *α*_*d*_. Low differentiation rates coupled with high proliferative capacity yield a high-ratio environment characterized by significant stochastic noise.

**Supplementary Video 1: Visual Simulation for HANG-AB3L with 5** *T*_1_ **NPAs at t=0**. The simulation was initiated with *α*_*pp*,0_ = *α*_*d*,0_ = 0.4. All other initial parameters were set as given in Table 2.

**Supplementary Video 2: Visual Simulation for HANG-AB3L with 100** *T*_1_ **NPAs at t=0**.The simulation was initiated with *α*_*pp*,0_ = 0.4, *α*_*d*,0_ = 0.3. All other initial parameters were set as given in Table 2.

**Supplementary Video 3: Visual Simulation for HANG-AB3L with 400** *T*_1_ **NPAs at t=0**.The simulation was initiated with *α*_*pp*,0_ = 0.4, *α*_*d*,0_ = 0.3. All other initial parameters were set as given in Table 2.

**Supplementary Video 4: Visual Simulation for HANG-AB3L with 800** *T*_1_ **NPAs at t=0**.The simulation was initiated with *α*_*pp*,0_ = 0.4, *α*_*d*,0_ = 0.3. All other initial parameters were set as given in Table 2.

## Acknowledgements

Authors would like to thank D. Tüfekçi for contributions on single layered HANG model; O. M. Anlı and U.Pamuk for inspiration and support for the manuscript.

## Declarations

The research is not funded by a third-party organization. Authors declare no conflict of interest. Codes for HANG-AB3L model will be available on Github upon publication (https://github.com/POZLAB/). The first draft of the manuscript was written by Pınar Ö z and all authors commented on previous versions of the manuscript. All authors read and approved the final manuscript.

## CRediT

Conceptualization: Pınar Ö z; Methodology: Pınar Ö z, Abdulsamet Atbaşı; Formal analysis and investigation: Pınar Ö z; Writing - original draft prepa-ration: Pınar Ö z; Writing - review and editing: Pınar Ö z, Abdulsamet Atbaşı; Supervision: Pınar Ö z.

## References

Aimone, J.B., Gage, F.H.: Modeling new neuron function: a history of using computational neuroscience to study adult neurogenesis. Eur J Neurosci 33(6), 1160–1169 (2011) 10.1111/j.1460-9568.2011.07615.x

Aimone, J.B.: Computational modeling of adult neurogenesis. Cold Spring Harb Perspect Biol. 8(4), 018960 (2016) 10.1101/cshperspect.a018960

Bast, L., Calzolari, F., Strasser, M.K., Hasenauer, J., Theis, F.J., Ninkovic, J., Marr, C.: Increasing neural stem cell division asymmetry and quiescence are predicted to contribute to the age-related decline in neurogenesis. Cell Rep 25(12), 3231–3240 (2018) 10.1016/j.celrep.2018.11.088

Barton, A., Fendrik, A.J., Rotondo, E.: A stochastic model of neurogenesis controlled by a single factor. J Theor Biol 355, 77–82 (2014) 10.1016/j.jtbi.2014.03.038

Baser, A., Skabkin, M., Kleber, S., Dang, Y., Gülcüler Balta, G.S., Kalamakis, G., et al.: Onset of differentiation is post-transcriptionally controlled in adult neural stem cells. Nature 566(7742), 100–104 (2019) 10.1038/s41586-019-0888-x

Bonaguidi, M.A., Wheeler, M.A., Shapiro, J.S., Stadel, R.P., Sun, G.J., Ming, G.-l., Song, H.: In vivo clonal analysis reveals self-renewing and multipotent adult neural stem cell characteristics. Cell 145(7), 1142–1155 (2011) 10.1016/j.cell.2011.05.024

Doetsch, F., Caille, I., Lim, D.A., García-Verdugo, J.M., Alvarez-Buylla, A.: Subventricular zone astrocytes are neural stem cells in the adult mammalian brain. Cell 97(6), 703–716 (1999) 10.1016/S0092-8674(00)80783-7

Encinas, J.M., Michurina, T.V., Peunova, N., Park, J.-H., Tordo, J., Peterson, D.A., et al.: Division-coupled astrocytic differentiation and age-related depletion of neural stem cells in the adult hippocampus. Cell Stem Cell 8(5), 566–579 (2011) 10.1016/j.stem.2011.03.010

Encinas, J.M., Sierra, A.: Neural stem cell deforestation as the main force driving the age-related decline in adult hippocampal neurogenesis. Behav Brain Res 227(2), 433–439 (2012) 10.1016/j.bbr.2011.10.010

Kuhn, H.G., Dickinson-Anson, H., Gage, F.H.: Neurogenesis in the dentate gyrus of the adult rat: age-related decrease of neuronal progenitor proliferation. J Neurosci 16(6), 2027–2033 (1996) 10.1523/JNEUROSCI.16-06-02027.1996

Kempermann, G., Song, H., Gage, F.H.: Neurogenesis in the adult hippocampus. Cold Spring Harb Perspect Biol 7(9), 018812 (2015) 10.1101/cshperspect.a018812

Li, B., Sierra, A., Deudero, J.J., Semerci, F., Laitman, A., Kimmel, M., Maletic-Savatic, M.: Multitype bellman-harris branching model provides biological predictors of early stages of adult hippocampal neurogenesis. BMC Syst Biol 11(Suppl 5), 90 (2017) 10.1186/s12918-017-0468-3

Lampada, A., Taylor, V.: Notch signaling as a master regulator of adult neurogenesis. Front Neurosci 17, 1179011 (2023) 10.3389/fnins.2023.1179011

Lehotzky, D., Zupanc, G.K.H.: Cellular automata modeling of stem-cell-driven development of tissue in the nervous system. Dev Neurobiol. 79(5), 497–517 (2019) 10.1002/dneu.22686

Míguez, D.G.: A branching process to characterize the dynamics of stem cell differentiation. Sci Rep 5(1), 13265 (2015) 10.1038/srep13265

Ming, G.-l., Song, H.: Adult neurogenesis in the mammalian brain: significant answers and significant questions. Neuron 70(4), 687–702 (2011) 10.1016/j.neuron.2011.05.001

Namba, T., Mochizuki, H., Suzuki, R., Onodera, M., Yamaguchi, M., Namiki, H., et al.: Time-lapse imaging reveals symmetric neurogenic cell division of gfap-expressing progenitors for expansion of postnatal dentate granule neurons. PLoS One 6(9), 25303 (2011) 10.1371/journal.pone.0025303

Obernier, K., Alvarez-Buylla, A.: Neural stem cells: origin, heterogeneity and regulation in the adult mammalian brain. Development 146(4), 156059 (2019) 10.1242/dev.156059

Obernier, K., Cebrian-Silla, A., Thomson, M., Parraguez, J.I., Anderson, R., Guinto, C., et al.: Adult neurogenesis is sustained by symmetric self-renewal and differentiation. Cell Stem Cell 22(2), 221–234 (2018) 10.1016/j.stem.2017.12.011

Öz, P.: A semi-stochastic numerical model of adult hippocampal neurogenesis. Süleyman Demirel Üniv Fen Bilim Enst Derg 23(1), 195–203 (2019) 10.19113/sdufenbed.471807

Pilz, G.-A., Bottes, S., Betizeau, M., Jörg, D.J., Carta, S., April, S., Simons, B.D., Helmchen, F., Jessberger, S.: Live imaging of neurogenesis in the adult mouse hippocampus. Science 359(6376), 658–662 (2018) 10.1126/science.aao5056

Postel, M., Karam, A., Pézeron, G., Schneider-Maunoury, S., Clément, F.: A multiscale mathematical model of cell dynamics during neurogenesis in the mouse cerebral cortex. BMC Bioinformatics 20(1), 470 (2019) 10.1186/s12859-019-3018-8

Stopka, A., Boareto, M.: A stochastic model of adult neurogenesis coupling cell cycle progression and differentiation. J Theor Biol 475, 60–72 (2019) 10.1016/j.jtbi.2019.05.014

Slater, J.L., Landman, K.A., Hughes, B.D., Shen, Q., Temple, S.: Cell lineage tree models of neurogenesis. J Theor Biol 256(2), 164–179 (2009) 10.1016/j.jtbi.2008.09.034

Silva-Vargas, V., Delgado, A.C., Doetsch, F.: Symmetric stem cell division at the heart of adult neurogenesis. Neuron 98(2), 246–248 (2018) 10.1016/j.neuron.2018.04.005

Ter Hoeven, E., Kwakkel, J., Hess, V., Pike, T., Wang, B., Kazil, J.: Mesa 3: agent-based modeling with Python in 2025. J Open Source Softw 10(107), 7668 (2025) 10.21105/joss.07668

Van Schepdael, A., Ashbourn, J.M.A., Beard, R., Miller, J., Geris, L.: Mechanisms of cell migration in the adult brain: modelling subventricular neurogenesis. Comput Methods Biomech Biomed Engin 16(10), 1096–1105 (2013) 10.1080/10255842.2013.773979

Wu, Y., Bottes, S., Fisch, R., Zehnder, C., Cole, J.D., Pilz, G.-A., Helmchen, F., Simons, B.D., Jessberger, S.: Chronic in vivo imaging defines age-dependent alterations of neurogenesis in the mouse hippocampus. Nat Aging 3(4), 380–390 (2023) 10.1038/s43587-023-00370-9

Ziebell, F., Dehler, S., Martin-Villalba, A., Marciniak-Czochra, A.: Revealing age-related changes of adult hippocampal neurogenesis using mathematical models. Development 145(1), 153544 (2018) 10.1242/dev.153544

Ziebell, F., Martin-Villalba, A., Marciniak-Czochra, A.: Mathematical modelling of adult hippocampal neurogenesis: effects of altered stem cell dynamics on cell counts and bromodeoxyuridine-labelled cells. J R Soc Interface 11(94), 20140144 (2014) 10.1098/rsif.2014.0144

