## Supplementary figures and images for "A Three-Layered Agent-Based Model of Adult Hippocampal Neurogenesis (HANG-AB3L) with Stochastic Cell Fate Determination"

### Supplementary Figure 1

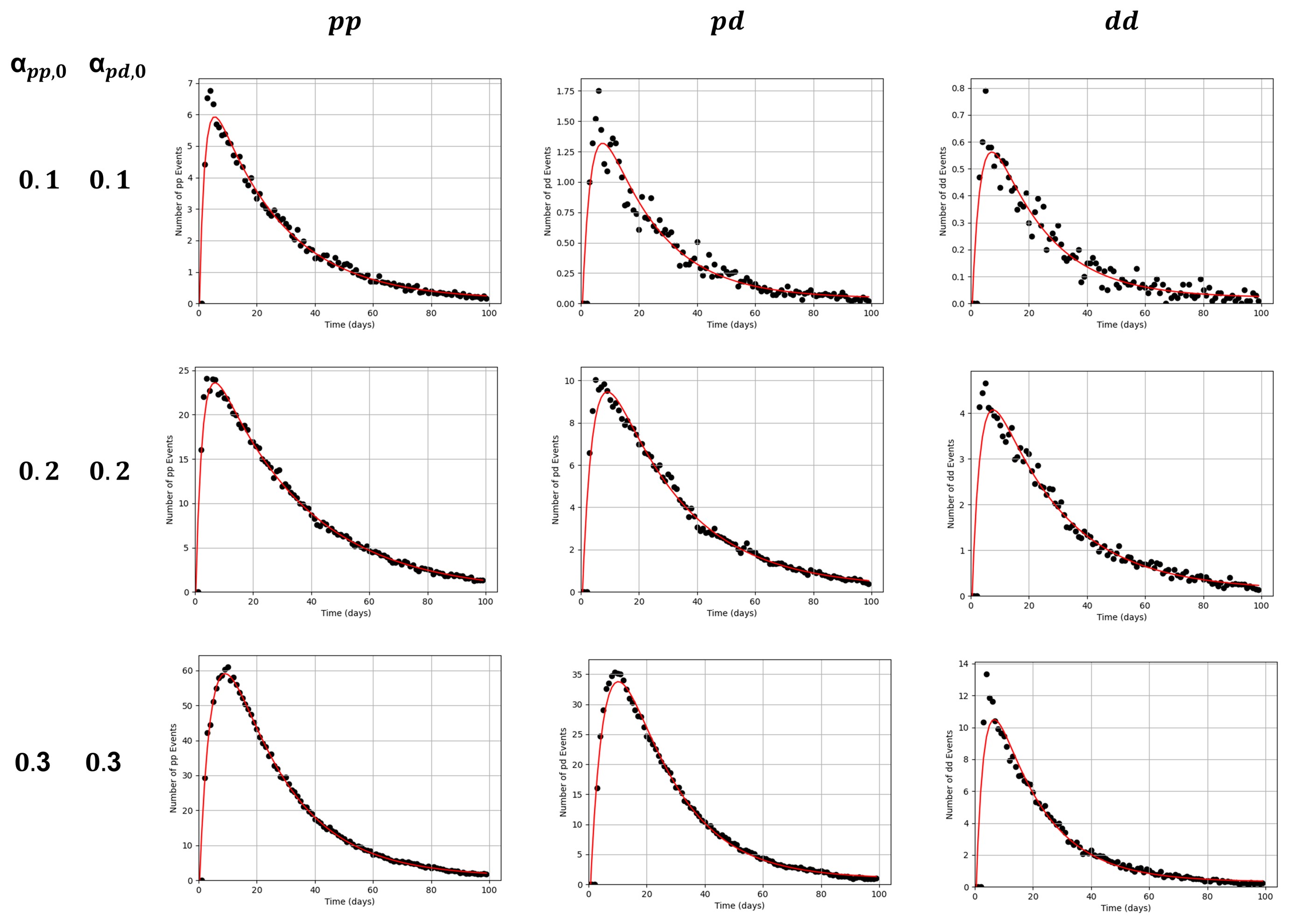

### Supplementary Figure 2

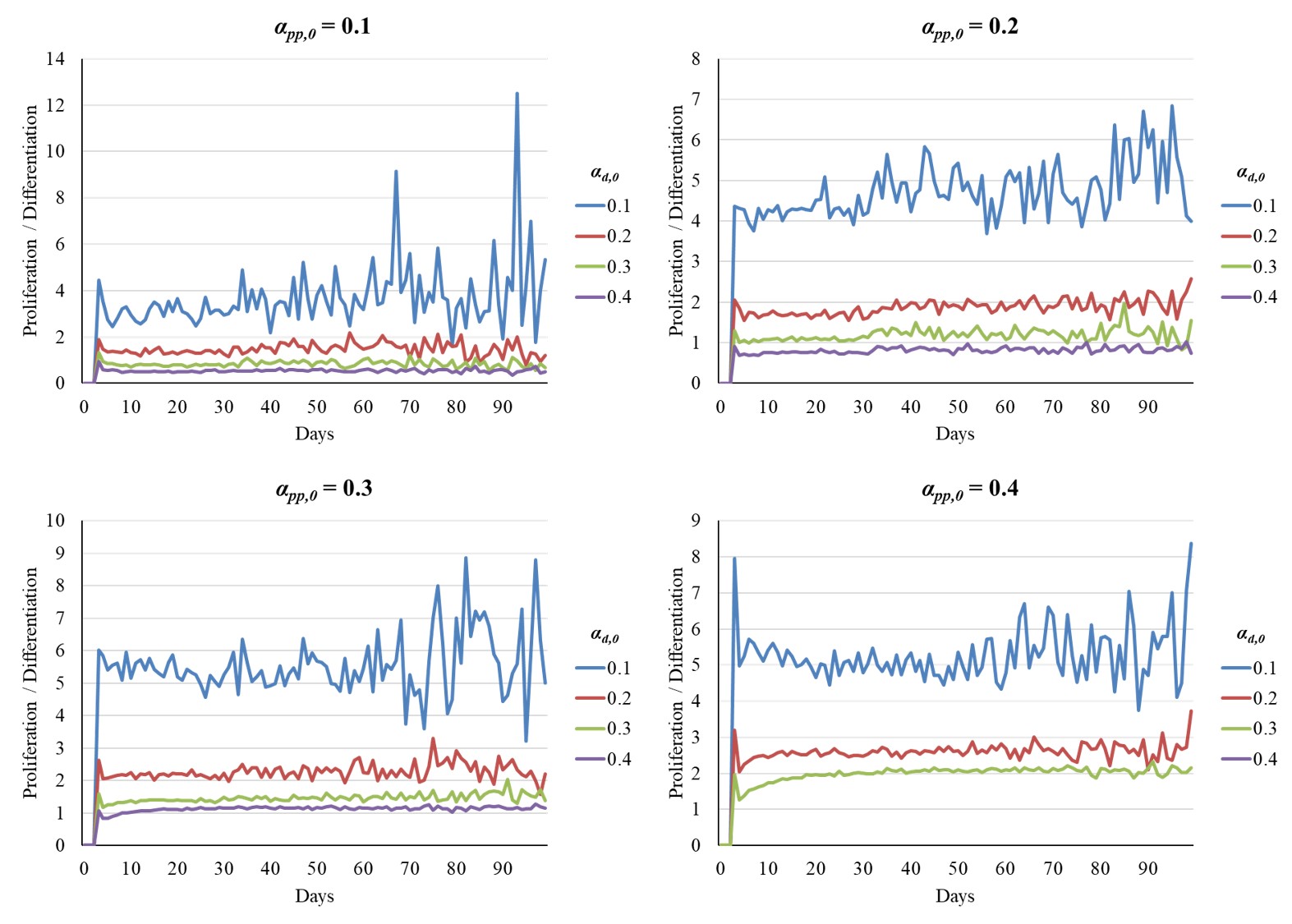

### Supplementary Figure 3

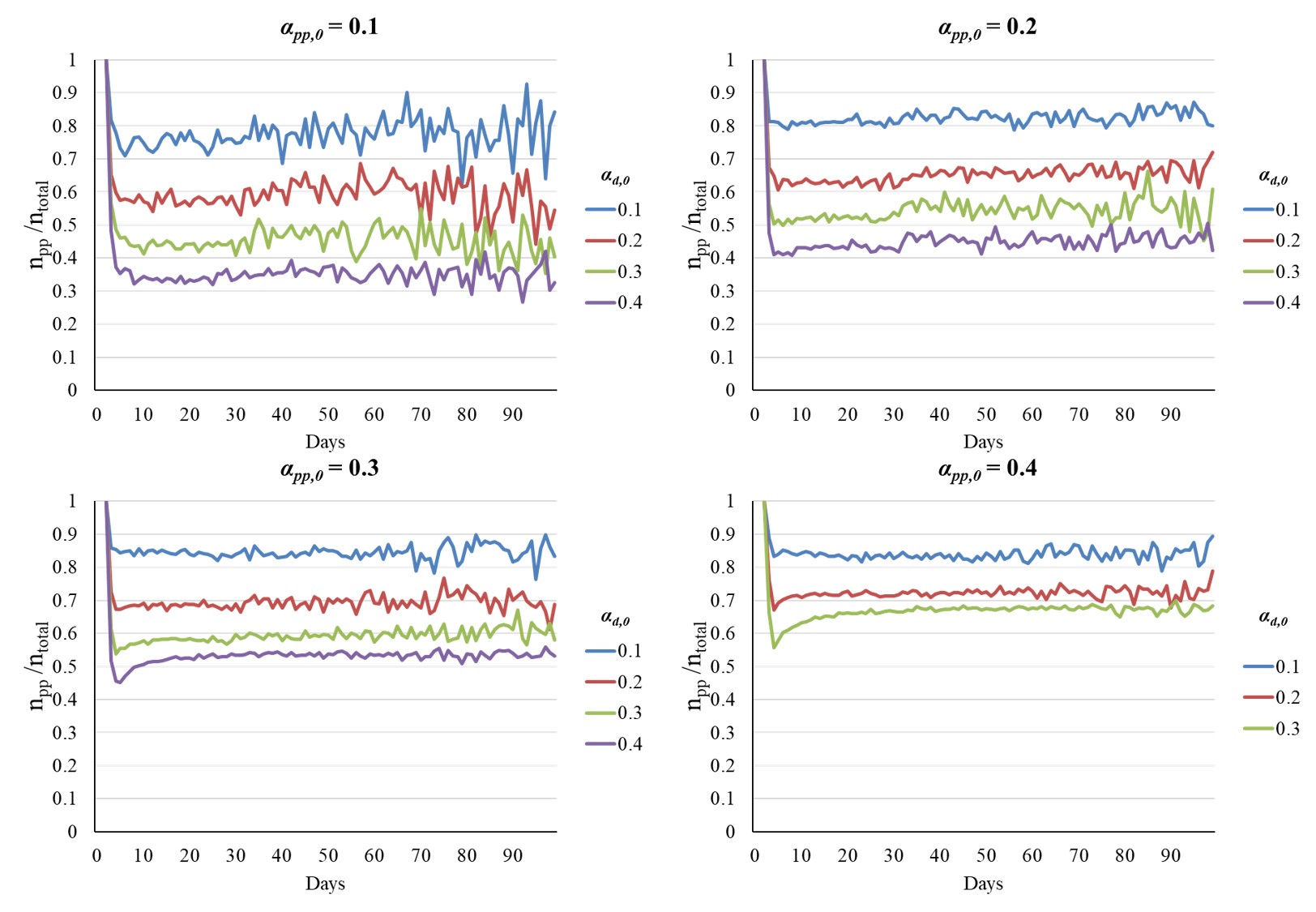
